# Tatton-Brown-Rahman Syndrome-associated *DNMT3A* mutations de-repress cortical interneuron differentiation to disrupt neuronal network function

**DOI:** 10.1101/2025.06.24.661324

**Authors:** Gareth Chapman, Julianna J. Determan, John R. Edwards, Faiza Batool, James E. Huettner, Ramachandran Prakasam, Sydney R. Crump, Yasmin Razia, Sofia Malik, Travis E. Law, Haley Jetter, Harrison W. Gabel, Kristen L. Kroll

## Abstract

Pathogenic mutations in DNMT3A cause Tatton-Brown-Rahman Syndrome (TBRS), a disorder characterized by somatic overgrowth of multiple tissues including the brain and intellectual disability (OGID). Here, we investigated TBRS etiology using new human pluripotent stem cell models, modeling varying levels of TBRS-associated loss of DNMT3A function. We identified lineage-specific overgrowth in TBRS ventral forebrain medial ganglionic eminence (MGE)-like progenitors, due in part to increased signaling through the PIK3/AKT/mTOR pathway that could be modulated to ameliorate this phenotype. By contrast, reduced DNA methylation during MGE-like progenitor differentiation into GABAergic interneurons caused premature expression of neuronal and synaptic genes, triggering precocious neuronal maturation. As a result, TBRS GABAergic neurons exhibited sufficient hyperactivity to alter the development and structure of neuronal networks, likely contributing to the intellectual disability and autism spectrum disorder common to TBRS patients. Together, this work elucidates new roles for DNMT3A-mediated gene repression in human cortical development, identifying critical requirements for regulating GABAergic neuron production and neuronal network function. These findings also support potential relationships between pathogenic mechanisms underlying TBRS and other OGIDs, including PIK3CA-related overgrowth syndrome and Weaver Syndrome, thus providing a foundation for future studies to identify common paradigms to treat these related disorders.

## Introduction

Overgrowth and intellectual disability disorders (OGIDs) are a group of rare neurodevelopmental disorders (Atterton et al. 2025) often caused by mutations either in proteins that mediate epigenetic regulation or in components of the PIK3/AKT/mTOR signaling cascade (Tatton-Brown et al. 2017). Characteristic of this, mutations in the *de novo* DNA methyltransferase *DNMT3A* cause Tatton-Brown-Rahman Syndrome (TBRS), a neurodevelopmental disorder involving autism spectrum disorder (ASD), brain overgrowth, and intellectual disability (ID) (Ostrowski and Tatton-Brown 1993; Tatton-Brown et al. 2018; Lane et al. 2020; Thomas et al. 2024). Several TBRS-associated mutations have been identified, each of which causes distinct levels of DNMT3A loss of function (Smith et al. 2021), with a majority involving heterozygous missense mutations in functional domains of *DNMT3A,* while other classes of mutation occur more rarely (Tatton-Brown et al. 2018).

Expression data from human postmortem (Kang et al. 2011) and mouse brain samples (Feng et al. 2005) show that DNMT3A expression begins during mid to late gestation, overtaking DNMT3B as the predominant *de novo* methyltransferase during early mouse neurodevelopment (Watanabe et al. 2006), where it regulates neural progenitor differentiation (Wu et al. 2012). However, despite the presence of DNMT3A during mouse brain development, TBRS mouse models exhibit anatomical and behavioral deficits reminiscent of TBRS with no evidence of brain overgrowth (Christian et al. 2020; Smith et al. 2021; Beard et al. 2023). Additionally, as these TBRS modelling efforts have focused on postnatal development to date (Christian et al. 2020; Smith et al. 2021; Beard et al. 2023), assessing how TBRS-associated loss of DNMT3A function alters early neurodevelopment to contribute to TBRS etiology remains critical. Furthermore, while the etiology of several other OGID disorders also remains understudied, particularly in human models, recent evidence supports a requirement for the OGID-associated gene *EZH2* in restraining human neuronal differentiation (Ciceri et al. 2024), while findings made in mouse models suggest that EZH2 and DNMT3A function are interrelated (Li et al. 2022). Therefore, investigating TBRS-associated cellular etiology and testing whether convergent molecular mechanisms contribute to other forms of OGID builds an essential foundation for developing both TBRS-specific and shared interventions across OGIDs.

Here we derived new pluripotent stem cell (hPSC) models of TBRS and studied how *DNMT3A* loss of function (LOF) mutations alter key aspects of human cortical development. TBRS models showed increased progenitor proliferation specific to medial ganglionic eminence (MGE)-like ventral forebrain neuronal progenitors (V-NPCs) and associated with increased signaling through the PIK3/AKT/mTOR pathway. TBRS models also exhibited increased neurogenesis and neuronal maturation in both 2-D and 3-D models of GABAergic interneuron development, correlating with precocious neuronal gene expression resulting from reduced DNA methylation. This resulted in GABAergic neuron hyperactivity sufficient to disrupt neuronal network synchrony. Together, this work demonstrates that GABAergic neurons are particularly sensitive to TBRS-associated DNMT3A mutation and also identifies convergent mechanisms that may be associated with pathogenic mutation of related OGID genes.

## Results

### TBRS models show increased V-NPC proliferation and mTOR signaling

To study the impact of TBRS-associated mutations on neuronal development and function we established four new sets of TBRS models spanning different levels of DNMT3A loss of function (LOF), consistent with the range of DNMT3A LOF observed across TBRS patient alleles (Nguyen et al. 2019; Christian et al. 2020; Smith et al. 2021; Beard et al. 2023). These included an induced pluripotent stem cell (iPSC) model from a patient with a p.R882H mutation (882), a recurrent severe loss of function (LOF) variant that we corrected to generate an isogenic control (C-WT, Fig. 1a), a human embryonic stem cell (hESC) model (WT) with variant knock-in of a p.P904L mutation (904), and an hESC model carrying small deletions near the end of the gene in both DNMT3A alleles and predicted to disrupt methyltransferase function (KO, Fig. 1b). We also derived two additional iPSC models from a patient with full gene deletion of one DNMT3A allele (Del1/2), which we paired with two unrelated sex matched control iPSC models from neurotypical individuals (Con1/2, Fig. 1c). Finally, we established two CRISPRi DNMT3A knockdown models (G1/G2) with their isogenic control (KRAB) (Fig. 1d). Total DNMT3A protein levels across these human pluripotent stem cell models (hPSCs) demonstrate little change in the 882 or 904 models, as expected (Fig. 1e-f), but significant loss of DNMT3A protein in the KO, Del1/2, and G1/G2 models (Fig. 1f-h).

**Fig. 1:**
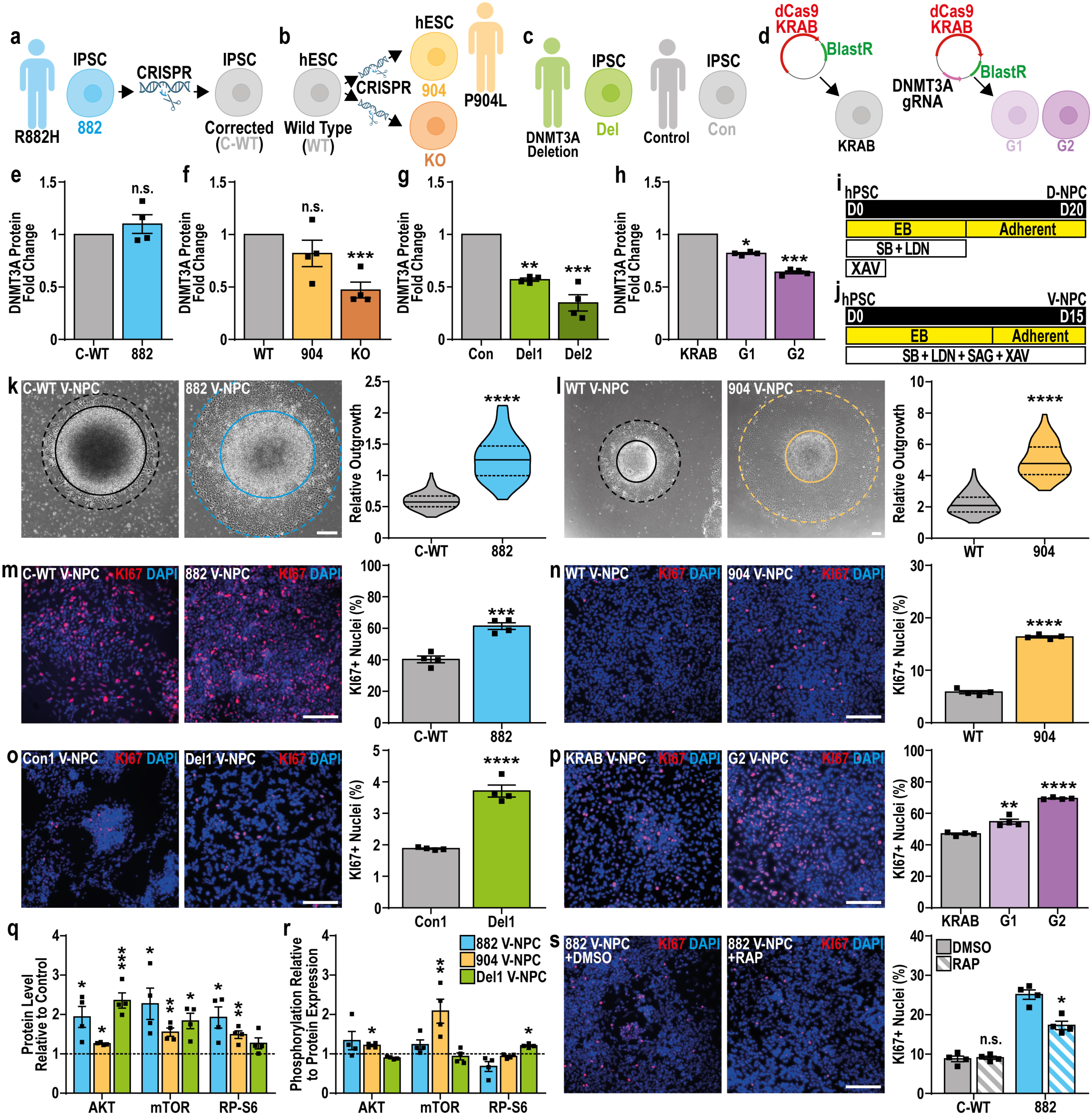
TBRS models show increased V-NPC proliferation and mTOR signaling. **a,** iPSCs carrying a p.R882H mutation in DNMT3A (882), generated from a TBRS patient and corrected to generate an isogenic control (C-WT). **b,** Control hESCs {WT), modified to mimic a TBRS patient with heterozygous p.P904Lmutation (904) via knock-in, which also created a model with small <10 bp mutations in both DNMT3Aalleles (KO). **c,** Two clonal iPSC lines were generated from a TBRS patient with heterozygous deletion of the DNMT3Agene (Del1/2) and compared with sex-matched unrelated control iPSCs (Con1/2). **d,** CRISPRi hPSC models were generated using two gRNAs targeting DNMT3A (G1/2) and compared with hPSCs without gRNA (KRAB). **e-h,** Quantification of DNMT3A protein expression in **(e)** 882, (f) 904, (f) KO, **(g)** Del1/2, and **{h)** G1/2 TBRS models versus matched controls.**i-j,** Schematic representation of (i) D-NPC specification and U) V-NPC specification. **k-I,** Representative images and quantification of relative neurosphere outgrowth {dotted line) from plated neurospheres {bold line) at D12 of **(k)** 822 and {I) 904 V-NPC specification versus matched controls. **m-p,** Representative images and quantification of Kl67 immunopositivity in **(m)** 822, **(n)** 904, **(o)** Del1 and **(p)** G1/2 V-NPCs versus matched controls. **q-r,** Quantification of relative **(q)** abundance and **(r)** phosphorylation of AKT, mTOR, and ribosomal protein S6 in TBRS V-NPCs (882, 904 and DeI1) versus matched controls {dotted line). **s,** Representative images and quantification of Kl67 positivity in 882 and C-WT V-NPCs after treatment with rapamycin (RAP) or vehicle control (DMSO). Data is represented as mean +/- SEM **(e-h,m-s)** or as distributions, with median {bold line) and upper and lower quartiles {dotted lines) indicated **(k-I),** and analyzed by Students t-test, versus matched controls **(e-h,k-s).** n=4 biological replicate experiments for all conditions, pValues: *p<0.05;**p<0.01;***p<0.001;****p<0.0001. Scale bars=200 µm **(k-I)** and 100 µm **(m-p,s).**

As brain overgrowth is central to TBRS clinical presentation, we first assessed whether these TBRS models exhibited altered growth during specification of neural progenitor cells (NPCs) with a dorsal (D-, Fig. 1i) or ventral (V-, Fig. 1j) telencephalic character. Initial observations highlighted an increase in neurosphere outgrowth in V-NPCs but not D-NPCs across the 882, 904 and Del1 models (Fig. 1k-l, Supplemental Fig. S1a-d). Further quantification of Ki67+ cells substantiated increased proliferation specific to V-NPCs across these TBRS models (Fig. 1m-o, Supplemental Fig. S1e-g) while assessments of lineage specific markers (PAX6 in D-NPCs or NKX2.1 in V-NPCs) showed no significant difference between control and TBRS NPCs (Supplemental Fig. S1h-m). Similar assessments in KO, Del2, and CRISPRi (G1/G2) V-NPCs also highlighted increased proliferation specific to V-NPCs (Fig. 1p, Supplemental Fig. S2a-d), solidifying this as a core TBRS-related phenotype.

Mutations in PIK3/AKT/mTOR pathway genes cause a related OGID disorder, PIK3CA-related overgrowth syndrome (PROS). This pathway is an established regulator of NPC proliferation (Romanyuk et al. 2024) and is related to DNMT3A activity in myeloid leukemia (Dai et al. 2017). Therefore, we next assessed if signaling through this pathway was altered in TBRS V-NPCs, finding that expression and/or phosphorylation of AKT3, mTOR and the mTOR target, ribosomal protein S6, increased specifically and consistently across TBRS V-NPCs (Fig. 1q-r, Supplemental Fig. S3a-b) while we observed no consistent changes in D-NPCs (Supplemental Fig. S3c-d). This confirmed that increased PIK3/AKT signaling was associated with increased V-NPC proliferation. We further tested whether reducing mTOR activity could correct TBRS-associated increases in V-NPC proliferation, demonstrating that rapamycin treatment significantly reduced V-NPC proliferation in TBRS models (Fig. 1s, Supplemental Fig. S3e-f). Together, these results highlight V-NPCs as highly sensitive to TBRS mutation, while also linking DNMT3A LOF with signaling disruptions underlying PROS, suggesting that convergent molecular mechanisms may underlie brain overgrowth across genetically distinct OGID disorders.

### DNMT3A restrains the onset of neuronal gene expression during V-NPC specification

To define molecular mechanisms disrupted by TBRS mutation in NPCs, we profiled gene expression changes between TBRS (882 and 904) and control V-NPCs, finding both distinct and shared effects across TBRS models (Fig. 2a-b, Supplemental Data 2). Focusing on differentially expressed genes (DEGs) that were similarly dysregulated in both models (shared-DEGs, Fig. 2b, Supplemental Data 2) we found the majority were up-regulated in TBRS models (Fig. 2b, Supplemental Data 2) and generally showed a greater effect size in 882 versus 904 V-NPCs (Supplemental Fig. S4a, Supplemental Data 2). While some of the shared-DEGs up-regulated in TBRS models encoded regulators of cell proliferation, consistent with the V-NPC overgrowth we observed, the majority were genes associated with neuronal differentiation (Fig. 2c, Supplemental Data 3), suggesting a second core function for DNMT3A. Assessing the expression of 30 of these shared-DEGs in 882 (severe LOF), 904 (mild LOF), G2 (mild LOF), and G1 (mildest LOF) V-NPCs, we found that increased expression of these genes correlated with DNMT3A LOF severity (Fig. 2d, Supplemental Fig. S4b), confirming a substantive role for DNMT3A in regulating both proliferative and neuronal gene expression. However, despite this substantive increase in neuronal gene expression, TBRS V-NPCs retained a progenitor identity (Supplemental Fig. S4c, Supplemental Data 2), signifying that DNMT3A LOF alone was insufficient to initiate neuronal differentiation. By contrast, parallel transcriptomic assessments of 882 D-NPCs revealed little effect of TBRS-mutation on gene expression, consistent with the lack of cellular phenotypes observed in TBRS D-NPCs (Supplemental Fig. S4d, Supplemental Data 2).

**Fig. 2:**
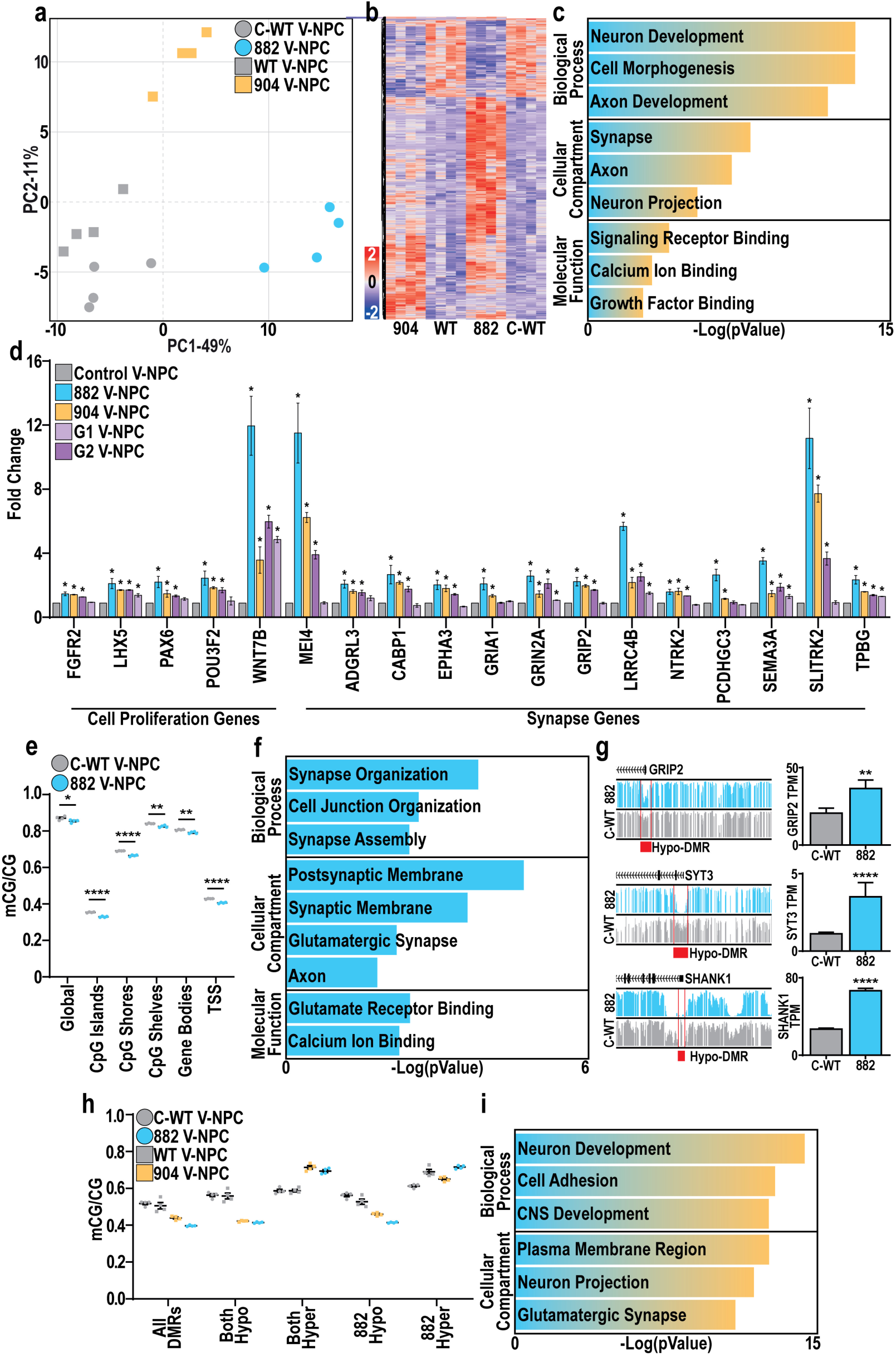
DNMT3A mediates epigenetic repression of pro-neuronal genes in V-NPCs. **a,** Principal component analysis of control (grey) and TBRS (882-blue and 904-orange) V-NPCs. **b,** Heatmap of shared DEGs across both 882 and 904 versus control (C-WT/WT) V-NPCs. **c,** Summary of GO enrichment analysis of DEGs upregulated in TBRS (882/904) versus control V-NPCs. **d,** Expression changes of genes associated with ‘cell proliferation’ and ‘synapse’ GO terms across TBRS (882, 904, G1/2) versus control V-NPCs. **e,** Changes in mCG/CG levels in C-WT versus 882 V-NPCs. **f,** Summary of GO enrichment analysis of 882 DEGs associated with an 882 hypo-DMR. **g,** Genomic browser views of mCG in C-WT and 882 V-NPCs, highlighting hypo-DMRs (red) associated with promoters of DEGs upregulated in 882 V-NPCs. **h,** mCG/CG differences at shared-DMRs (All), qualified by directionality of DMRs in both TBRS models (both-hyper or both-hypo) or specifically in the 882 model (882-hyper or 882-hypo). i, Summary of GO enrichment analysis of DEGs upregulated in TBRS (904 and 882) V-NPCs and associated with shared hypo-DMRs.Data is represented as mean +/-SEM **(d,e,g,h)** with individual biological replicates indicated **(e,h).** Data was analyzed by Student’s t-test versus isogenic controls **(d,e)** or by differential expression analysis **(g).** n=4 biological replicate experiments for all conditions, pValues: *p<0.05 **(d)** and *p<0.05; **p<0.01; ****p<0.0001**(e,g).**

We next investigated the proximal cause of the TBRS-associated transcriptomic dysregulation by identifying changes in DNA methylation (mCG) in 882 and 904 V-NPCs, identifying significant global and regional mCG decreases specific to 882 V-NPCs (Fig. 2e; Supplemental Fig. S5a). Of note, while DNMT3A also mediates non-CpG methylation in postnatal and adult mice (Beard et al. 2023), our hPSC-derived cultures do not exhibit appreciable levels of non-CpG methylation owing to their relative immaturity. Focusing on the 882 model, we identified differentially methylated regions (DMRs), most of which lost mCG in 882 V-NPCs (hypo-DMRs, Supplemental Data 4), with these sites commonly associated with synaptic and neuronal genes upregulated in TBRS V-NPCs (Fig. 2f-g, Supplemental Data 5). Examining mCG changes in both 882 and 904 V-NPCs, we found that 882 V-NPCs exhibited greater mCG loss than 904 V-NPCs across the superset of all DMRs present in both TBRS models (All DMRs, Fig. 2h) and at 882 hypo-DMRs (Fig. 2h), consistent with differential DNMT3A LOF across these models. When examining the subset of DMRs consistently hypo-methylated across both TBRS models, we found a similar reduction in mCG levels across both models (Fig. 2h). Furthermore, these shared-DMRs were often located at gene promoters (Supplemental Fig. S5b) of neuronal genes (Fig. 2i, Supplemental Data 5), many of which were also shared-DEGs (Supplemental Data 5). Similar analyses of 882 D-NPCs revealed substantive changes in mCG (Supplemental Data 4), but these changes did not correlate with substantive transcriptomic changes (Supplemental Data 2), indicating that TBRS-associated mutations do not grossly impact gene expression in D-NPCs. Together, these results demonstrate that TBRS-associated mutations disrupt the ability of DNMT3A to restrain both proliferation-associated gene expression and onset of neuronal gene expression in V-NPCs.

### TBRS organoids modeling ventral forebrain development exhibit increased neurogenesis

To examine the consequences of TBRS-associated alterations on NPC development, we profiled changes in the development of dorsally (D-ORG, Fig. 3a) and ventrally (V-ORG, Fig. 3b) patterned forebrain-like organoids. Drawing comparisons with the increased proliferation observed in our 2-D V-NPC derivation scheme, we found that the proliferative Ki67+ cell fraction likewise increased in TBRS (882, 904, and Del1) V-ORGs (Fig. 3c, Supplementary Fig. S6a), while assessing the V-NPC markers DLX2 and NKX2.1 revealed no consistent alterations in TBRS V-ORG specification (Supplementary Fig. S6b-c). Given the increased neuronal gene expression observed in TBRS V-NPCs, we also examined altered neurogenesis in TBRS V-ORGs, finding a significant increase in the TUJ1-immunopositive area (Fig. 3d-e), which was not due to loss of SOX2+ progenitors (Fig. 3d, f-g). Therefore, to assess whether this increase in TUJ1-immunopositive area resulted from increased neurogenesis, we also examined the proportion of cells positive for the pro-neurogenic marker ASCL1 or immature GABAergic neuron marker SST, finding an increase in both ASCL1+ and SST+ cells in TBRS V-ORGs (Fig. 3h-i, Supplementary Fig. S6d-e), substantiating an increased neurogenesis in TBRS V-ORGs.

**Fig. 3:**
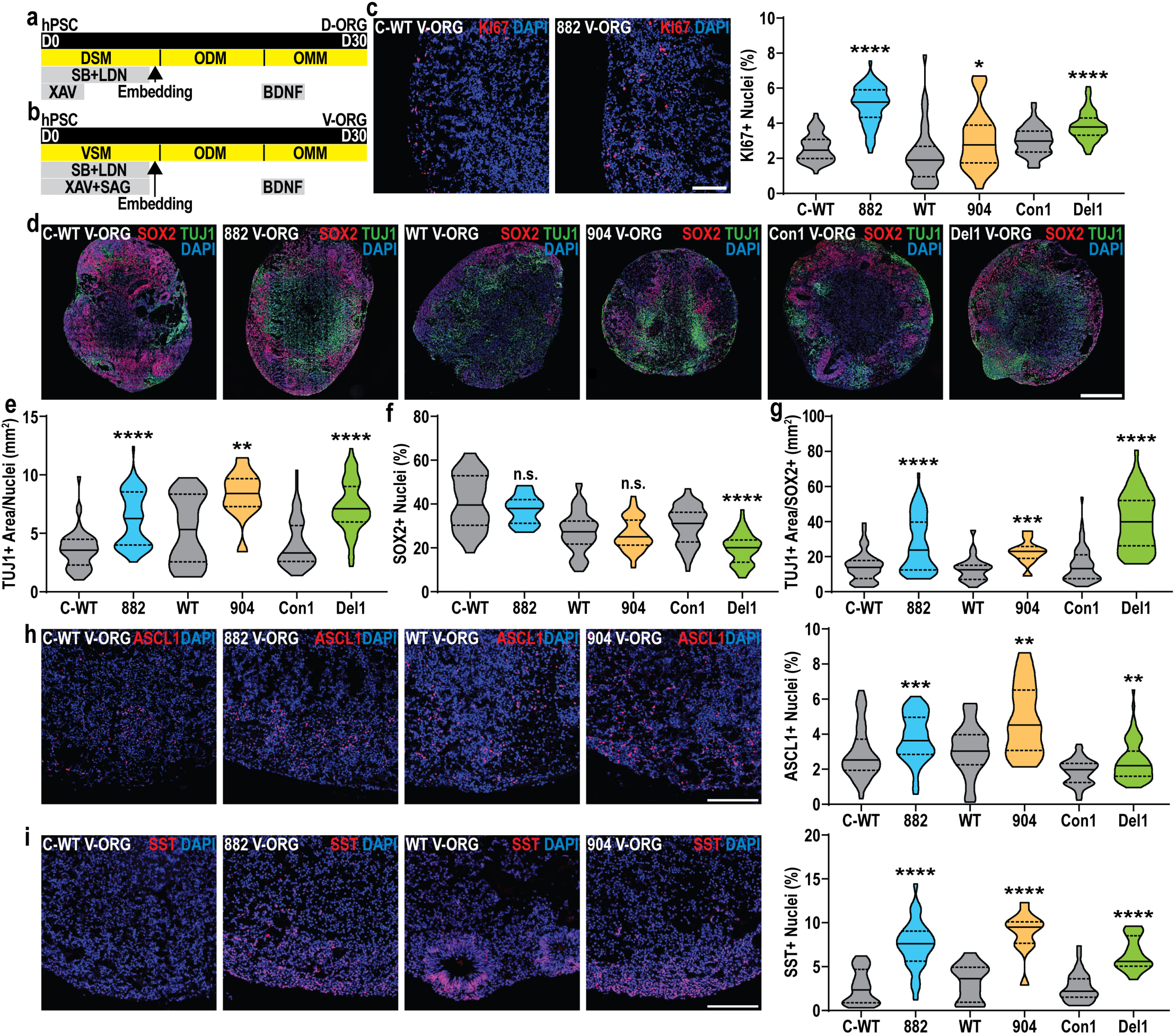
TBRS V-ORGs show increased proliferation and neurogenesis. a-b, Schematic representation of **(a)** D-ORG or **(b)** V-ORG differentiation. **c,** Representative images and quantification of Kl67+ cell fraction across TBRS and control V-ORGs.**d-g,** Representative images of **(d)** SOX2 and TUJ1 staining in TBRS and control V-ORGs, alongside quantification of **(e)** TUJ1+ area,(f) SOX2+ fraction (% of DAPI+ nuclei) and **(g)** TUJ1+ area normalized to SOX2+ cells. **h-i,** Representative images and quantification of the **(h)** ASCL1+ and (i) SST+ cell fraction across TBRS and control V-ORGs. All data is represented as distributions with median (bold line) and upper and lower quartiles (dotted lines) indicated and was analyzed by Mann-Whitney tests, comparing each TBRS model to its matched control. A minimum of 3 batches of organoids were prepared for each condition, with 3-9 organoids assessed per batch. pValues:*p<0.05; **p<0.01; ***p<0.001; ****p<0.0001;n.s.-non-significant. Scale bars=100 µm **(c,h-i)** and 500 µm (d).

Performing similar assessments in D-ORGs, we found no consistent evidence for altered proliferation (Supplemental Fig. S7a), again congruent with our prior observations in 2-D models. However, we did observe an increased proportion of SOX2+ cells in 904 D-ORGs with no significant change in the TUJ1 immunopositive region across TBRS D-ORGs (Supplementary Fig. S7b-c). Finally, we assessed changes in neurogenesis using ASCL1, the radial glial cell marker TBR2, and the early born cortical glutamatergic neuron marker TBR1, finding no consistent differences in the proportion of any of these populations in TBRS D-ORGs (Supplementary Fig. S7d-f). Therefore, consistent with our findings in 2-D models of NPC specification, our organoid models also show that TBRS-associated DNMT3A LOF has little effect on D-NPC development, while causing increased proliferation and neurogenesis during V-NPC development, suggestive of increased production and/or maturation of GABAergic neurons.

### Repressive H3K27me3 compensates for severe loss of DNA methylation

Prior work has also demonstrated a role for *EZH2* in restraining neuronal maturation during human glutamatergic neuron differentiation, of particular interest as pathogenic *EZH2* mutation can cause the OGID Weaver syndrome (Gibson et al. 2012; Tatton-Brown et al. 2013; Ciceri et al. 2024). As this effect was reminiscent of our findings for DNMT3A LOF mutation, we investigated if levels of histone H3 lysine 27 tri-methylation (H3K27me3) deposited by EZH2 were altered in our TBRS models. Identifying sites of differential H3K27me3 enrichment (DKSs) in both 882 D-NPCs and V-NPCs, we found that the majority gained H3K27me3 in the TBRS model (hyper-DKS, Fig. 4a-b, Supplemental Data 6). Interestingly, while D-NPC and V-NPC hyper-DKSs that intersected with 882 hypo-DMRs were associated with genes involved in early neural development (Fig. 4c-d, Supplemental Data 7), few of these were TBRS DEGs. Together, these data suggest that H3K27me3-mediated repression may compensate for mCG reduction resulting from TBRS-associated mutation, an interrelationship supported by the postnatal gain of H3K27me3 following the conditional knockout of *Dnmt3a* in mice (Li et al. 2022). Thus, we hypothesized that compensatory H3K27me3-mediated repression may occur in 882 NPCs to prevent transcriptomic dysregulation resulting from DNMT3A LOF mutation.

**Fig. 4:**
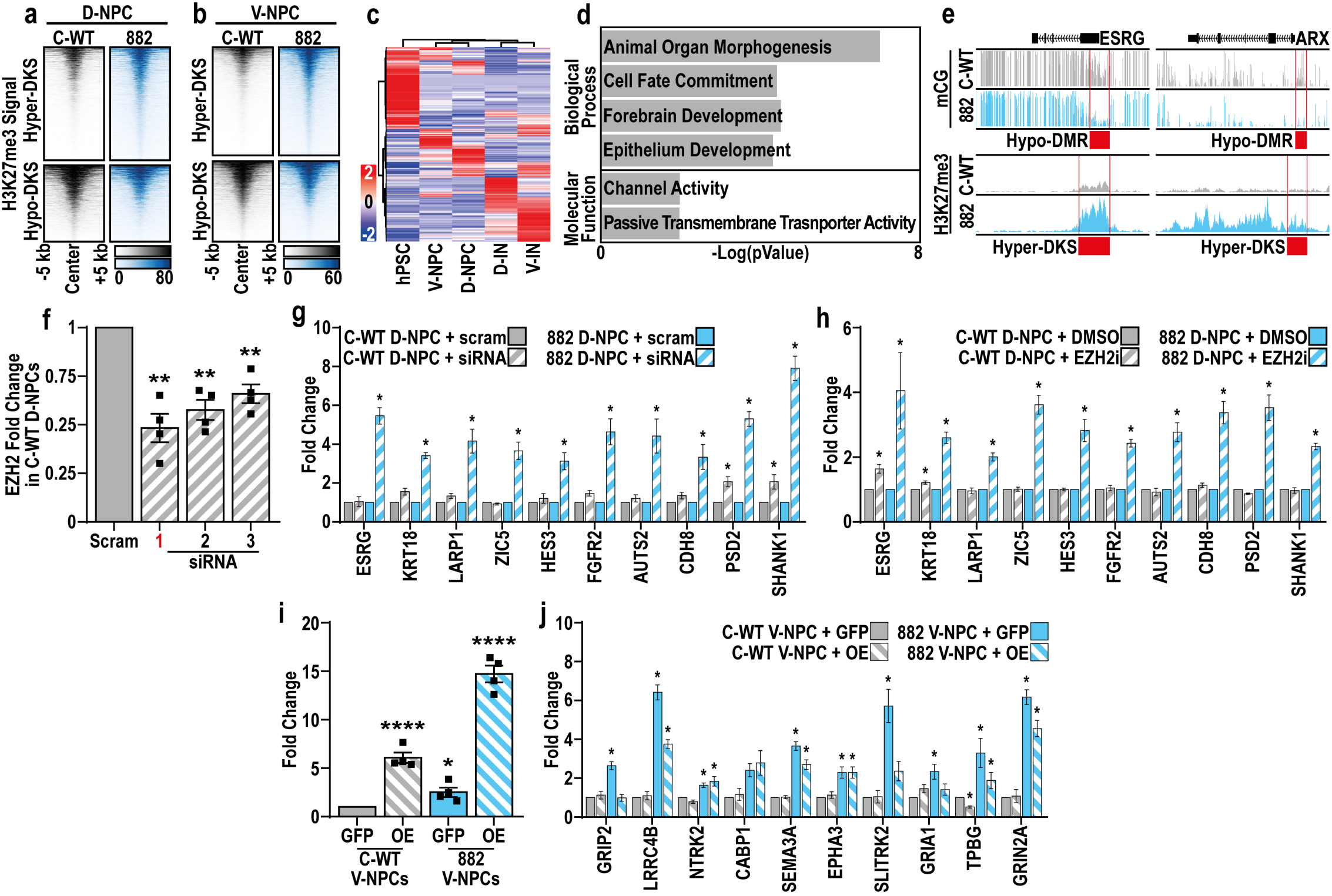
Increased H3K27me3 compensates for loss of DNA methylation specifically in R882H NPCs. a-b, H3K27me3 peaks with significantly increased (hyper-OKS) or decreased (hypo-OKS) enrichment in 882 versus control **(a)** D-NPCs or **(b)** V-NPCs.**c,** Expression of genes associated with hypo-DMRs and hypo-DKSs in 882 NPCs (D and V) across neuronal differentiations. **d,** Summary of GO enrichment analysis of non-DEGs associated with hypo-DMRs and hypo-DKSs in 882 NPCs (D and V). **e,** Genomic browser views of mCG and H3K27me3 in 882 and control V-NPCs, highlighting (red) hypo-DMRs and hyper-DKSs associated with promotors of two genes. **f,** Quantification of EZH2 mRNA levels in C-WT D-NPCs after treatment with either scrambled siRNA (scram) or siRNAs targeting EZH2 (1-3), highlighting selection of siRNA 1 for further study (red). **g-h,** Quantification of expression changes in 10 genes associated with 882 D-NPC hypo-DMRs and hyper-DKSs, after treatment with **(g)** siRNAs or **(h)** an EZH2-specific inhibitor. i, Quantification of EZH2 mRNA levels in C-WT and 882 V-NPCs after lentiviral transduction with a construct overexpressing either EZH2 (OE) or GFP (GFP). **j,** Quantification of expression changes in 10 genes upregulated in 882 V-NPCs under OE or GFP conditions. Data represented as mean+/-SEM**(f-j)** and was analyzed by one-way ANOVA, with comparison of treatment versus **(f-g)** scram, **(h)** DMSO or **(i-j)** GFP control treated conditions. n=4 biological replicate experiments for all conditions, pValues: *p<0.05; **p<0.01; ****p<0.0001**(f,i)** or *p<0.05 **(g-h,j).**

To test this hypothesis, we identified a set of genes in 882 D-NPCs that were associated both with hypo-DMRs and hyper-DKSs but were not TBRS DEGs (e.g. ESRG and ARX, Fig. 4e) and tested whether disrupting H3K27me3 deposition altered their expression. Either EZH2 knockdown in 882 TBRS D-NPCs using siRNAs (Fig. 4f, Supplemental Fig. S8a) or disruption of EZH2 activity with a selective chemical inhibitor (EZH2i) disrupted this compensatory repression, leading to increased target gene expression relative to controls (Fig. 4g-h). Significantly, these effects were not observed in D-NPCs with less severe DNMT3A LOF (904; Supplemental Fig. S8b), suggesting that compensatory H3K27me3 increase may be triggered by the more severe mCG loss in the 882 model. Given these findings, we examined whether the overexpression (OE) of EZH2 could reverse the gene dysregulation associated with loss of DNMT3A-mediated repression in the more severely impacted V-NPCs, finding that EZH2 OE partially rescued this TBRS-associated transcriptomic dysregulation (Fig. 4i-j). Together, these results indicate that repressive H3K27me3 compensates for reduced DNA methylation stemming from severe mCG loss, while increased EZH2 activity can ameliorate some TBRS-associated molecular phenotypes. These data support interrelated mechanisms of altered gene regulation underlying TBRS and other OGIDs such as Weaver syndrome resulting from pathogenic EZH2 mutation.

### TBRS mutations cause precocious maturation of GABAergic neurons

We next investigated whether the elevated neuronal gene expression found in TBRS V-NPCs persisted upon their differentiation into immature GABAergic cortical interneurons (V-INs; Fig. 5a). Reminiscent of our V-NPC analysis, we identified greater transcriptomic differences in 882 than 904 V-INs (Fig. 5b, Supplemental Data 8), while shared-DEGs upregulated in TBRS V-INs were again enriched for synaptic genes (Fig. 5c, Supplemental Data 9). Effect sizes of TBRS V-IN shared-DEGs were likewise generally larger in 882 versus 904 V-INs, as exemplified by 42 synapse-associated shared-DEGs (Fig. 5d, Supplemental Data 9). We also performed comparative gene set enrichment analysis with published data from the adult cortex of a TBRS mouse model (p.R878H) analogous to the 882 human mutation (Beard et al. 2023), finding shared upregulation of neuronal and synaptic genes (Supplemental Data 10). Together, these results highlight substantive evidence for synaptic gene dysregulation as a core phenotype associated with TBRS mutation.

**Fig 5.**
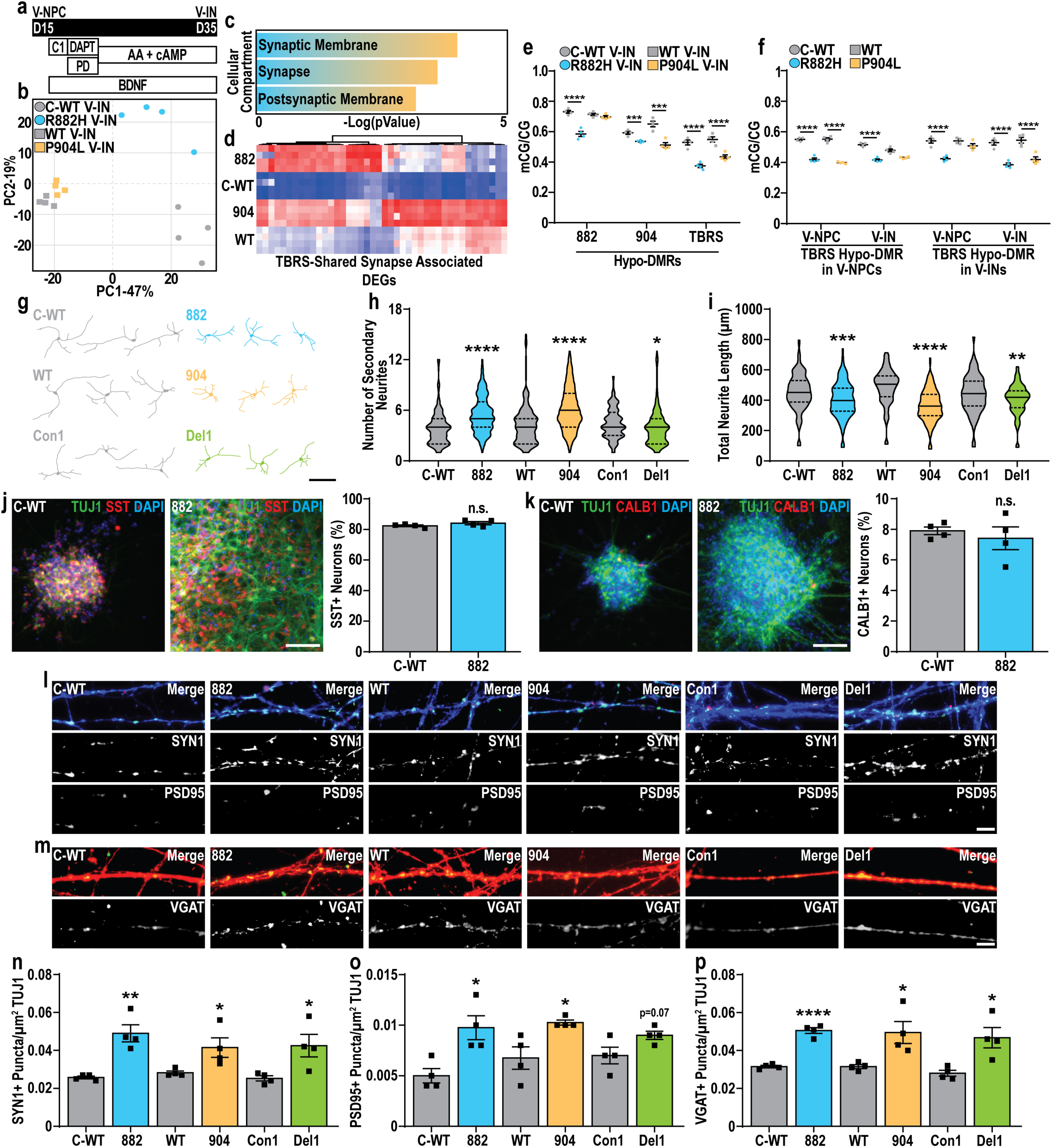
DNMT3A constrains neuronal maturation during interneuron differentiation. **a,** Experimental paradigm for generating immature GABAergic neurons (V-INs) from ventrally patterned neuronal progenitors (V-NPCs). **b,** Principal component analysis of control (grey) and TBRS (882-blue and 904-orange) V-INs.**c,** Summary of GO enrichment analysis of DEGs upregulated in TBRS V-INs.**d,** Heatmap of genes associated with the ‘synapse’ GO term and upregulated in TBRS V-INs. **e,** Changes in mCG/CG at 882, 904, or shared (TBRS) V-IN hypo-DMRs and matched control V-INs. **f,** Changes in mCG/CG at TBRS-shared hypo-DMRs identified in V-NPCs or V-INs across both TBRS and control V-NPCs and V-INs. **g,** Representative example traces of control and TBRS D30 V-INs used to quantify neuronal morphology. **h-i,** Quantification of neuronal morphology in control and TBRS D30 V-INs,including **(h)** number of secondary neurites and (i) total neurite length. **j-k,** Representative images and quantification of the proportion of U) SST or **(k)** CALB1 positive neurons in day (D) 40 C-WT or 882 V-IN cultures. **1-p,** Representative images and quantification of pre-synaptic marker **(l,n)** SYN1 and postsynaptic marker **(1,o)** PSD95 puncta density, and **(m,p)** VGAT, across TBRS and control D50 V-INs. Data was analyzed by Students I-test versus matched controls **(e,f,h,i,n-p).** n=4 biological replicate experiments for all conditions with a minimum of 20 neurons quantified for morphological measures per biological replicate,pValues: *p<0.05;**p<0.01;***p<0.001;****p<0.0001. Scale bars=100 µm **(g),** 50 µm **U,k)** and 10 µm **(1,m).**

To establish if continued upregulation of these synaptic genes resulted from persistent mCG loss in TBRS V-INs, we examined DNA-methylation in 882 and 904 V-INs, highlighting milder mCG losses than those observed in V-NPCs (Supplemental Fig. S9a-b). However, identifying DMRs across TBRS V-INs highlighted many hypo-DMRs (Supplemental Data 11), with 904 V-IN hypo-DMRs frequently detected in 882 V-INs, but not vice versa (Fig. 5e, Supplemental Data 11). Focusing on hypo-DMRs shared across 882 and 904 V-INs (Fig. 5f, Supplemental Data 11) and associating these with V-IN shared-DEGs again revealed synaptic gene enrichment (Supplemental Data 9). We also found that, despite their similar association with synaptic genes, 882 hypo-DMRs persisted through development at the same sites, while locations of 904 hypo-DMRs were more temporally specific (Fig. 5f, Supplemental Data 11). Together, these findings support persistently diminished DNA methylation and concomitant upregulation of synaptic genes as a continued feature during TBRS GABAergic neuron differentiation.

To assess how these molecular phenotypes affected neuronal maturation, we next assessed whether TBRS GABAergic neurons exhibited altered morphology, scoring control and TBRS (882, 904, and Del1) V-INs at D30. This analysis revealed an increase in ramification, specifically involving increased secondary neurite production at the expensive of overall neurite length in TBRS V-INs (Fig. 5g-i, Supplemental Fig. S10a), while also highlighting an increase in soma size of TBRS V-INs (Supplemental Fig. S10b). We also assessed whether cortical interneuron identity was disrupted by DNMT3A LOF by examining the production of CALB1+ and SST+ V-INs at day (D) 40, finding no significant difference in the fraction immunopositive for either marker in TBRS V-IN cultures (Fig. 5j-k, Supplemental Fig. S10c-f). Finally, we evaluated if synapse formation was disrupted in TBRS V-INs, finding increased synaptic marker density (SYN1, PSD95, and VGAT) in D50 TBRS V-INs (Fig. 5l-p). Together, these results confirm that TBRS-associated transcriptomic dysregulation during V-IN differentiation also causes premature V-IN maturation.

Performing parallel transcriptomic analysis in immature glutamatergic neurons (D-INs), we found substantive transcriptomic differences between TBRS (882 and 904) and control D-INs (Supplemental Fig. S11a-b), characterized by persistent expression of genes normally expressed in hPSCs and D-NPCs (Supplemental Fig. S11c-d, Supplemental Data 12-13). We tested whether this phenomenon was a consequence of altered development by bypassing NPC specification, instead deriving 882 and 904 glutamatergic-like neurons by inducible NGN2 overexpression (iGluts, Supplemental Fig. S11e); this revealed similar transcriptomic dysregulation across TBRS D-INs and iGluts (Supplemental Fig. S11f, Supplemental Data 12-13). We also found similar global losses of mCG across 882 but not 904 D-INs and iGluts (Supplemental Fig. S11g-j), while identifying DMRs in TBRS highlighted similar losses of mCG across D-INs and iGluts (Supplemental Fig. S11k; Supplemental Data 14). TBRS-shared hypo-DMRs in D-INs were often associated with neuronal and synaptic genes (Supplemental Data 13) and, consistent with this observation, we found that DMRs across 882 D- and V-INs showed similar mCG losses (Supplemental Fig. S11l, Supplemental Data 13-14). Together, our work above shows that TBRS mutations most profoundly alter GABAergic neuron development, due to persistent dysregulation of neuronal and synaptic gene expression, while TBRS glutamatergic neurons exhibit distinct transcriptomic changes and patterns of mCG loss, changes largely decoupled from altered NPC specification.

### TBRS associated GABAergic neuron dysfunction causes neuronal network hypersynchrony

We next assessed the functional consequences of TBRS mutation by performing patch clamp electrophysiology on glutamatergic or GABAergic neurons (matured D-INs or V-INs, respectively) plated in monoculture on rat astrocytes and further matured. Initial assessments showed no significant functional alterations in 882 glutamatergic neurons, consistent with the lack of cellular phenotypes during D-IN differentiation (Supplemental Fig. S12a-d, Supplemental Data 15). By contrast, 882 GABAergic neurons exhibited significant hyperactivity upon stimulation (Fig. 6a-b), without significant alterations to neuronal health (Fig. 6c, Supplemental Fig. S13a-d, Supplemental Data 16). This hyperactivity was characterized by an increased action potential (AP) amplitude and decreased AP halfwidth (Fig. 6d-e). We also observed increased sodium uptake, transient outward currents (TOC), and GABA responsiveness in 882 GABAergic neurons (Fig. 6f-h, Supplemental Fig. S13e). When similar data was generated for 904 glutamatergic or GABAergic neurons, combined, and analyzed, we likewise found evidence for neuronal hyperactivity upon stimulation (example in Fig. 6i, combined data in Fig. 6j), while neither neuronal cell type showed significant changes in other measures of neuronal health or activity (Supplemental Fig. S13f-k, Supplemental Data 15-16).

**Fig 6.**
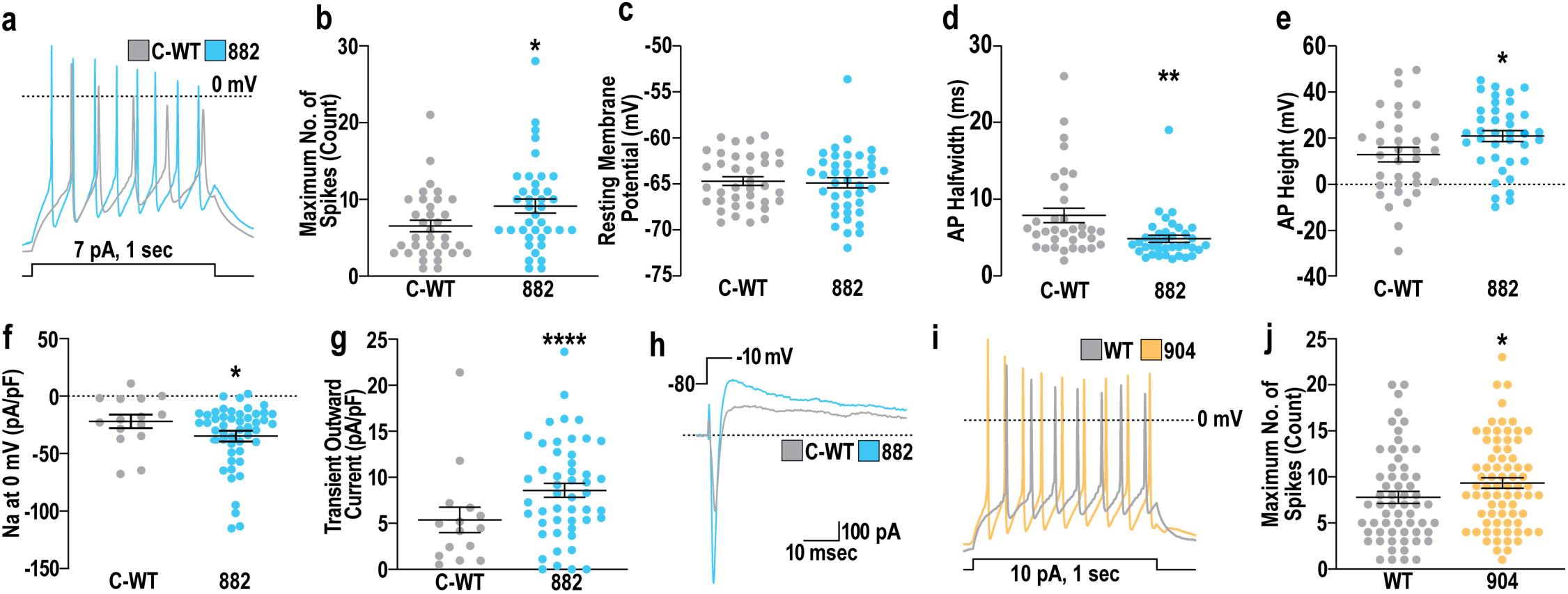
TBRS-associated mutations in DNMT3A cause GABAergic neuron hyperactivity. **a,** Representative example traces of action potentials in C-WT (grey) or 882 (blue) GABAergic neurons induced by 1 second of stimulation. **b,** Quantification of maximum spike number across C-WT and 882 GABAergic neurons during 1 second of stimulation. **c,** Resting membrane potential of C-WT and 882 GABAergic neurons. **d-e,** Quantification of action potential (AP) characteristics, including **(d)** AP halfwidth and **(e)** AP height of C-WT and 882 GABAergic neurons. **f-g,** Measurements of (f) sodium (Na) current density at 0 mV and **(g)** transient outward current after voltage step, both normalized by capacitance (pA/pf}.**h,** Example traces of current changes in C-WT and 882 GABAergic neurons during voltage step from -80 to -10 mV. i, Representative example traces of action potentials in WT (grey) or 904 (orange) GABAergic neurons induced by 1 second of stimulation. **j,** Quantification of maximum spike number, measured in both glutamatergic and GABAergic neurons (combined data analysis), during 1 second of stimulation. Significance was calculated by Rank Sum Test and results are presented as mean+/-SEM with data points representing measurements taken from individual neurons, *pValue<0.05;**pValue<0.01;***pValue<0.001.

We next used low density (LD) multi-electrode arrays (MEAs) to assess changes in neuronal network activity in co-cultures of control or TBRS glutamatergic and GABAergic neurons. This analysis showed divergent results across models, with 882 cultures showing increased activity (Fig. 7a-d; Supplemental Fig. S14a-b), while trends from 904 cultures were inconsistent with this finding (Supplemental Fig. S14c-g). Therefore, to study these phenotypes at greater resolution, we assessed neuronal network activity using high density (HD) MEAs, focusing on TBRS GABAergic neuron dysfunction. Examining TBRS or control day (D) 50 GABAergic neurons co-cultured with control iGluts, we observed no differences in mean firing rate (MFR) but increased numbers of electrodes exhibiting bursting behavior (Fig. 7e-g). Furthermore, in assessing the same cultures at D82, co-cultures containing TBRS GABAergic neurons had an increased MFR, with a greater effect size in cultures containing 882 versus 904 GABAergic neurons (Fig. 7h-i). This was coupled with a decreased inter-spike interval (ISI) and increased coefficient of variation (CoV) of ISI associated with both the 882 and 904 models, with cultures containing GABAergic 882 neurons again showing greater effect sizes (Supplemental Fig. S15a-b).

**Fig 7.**
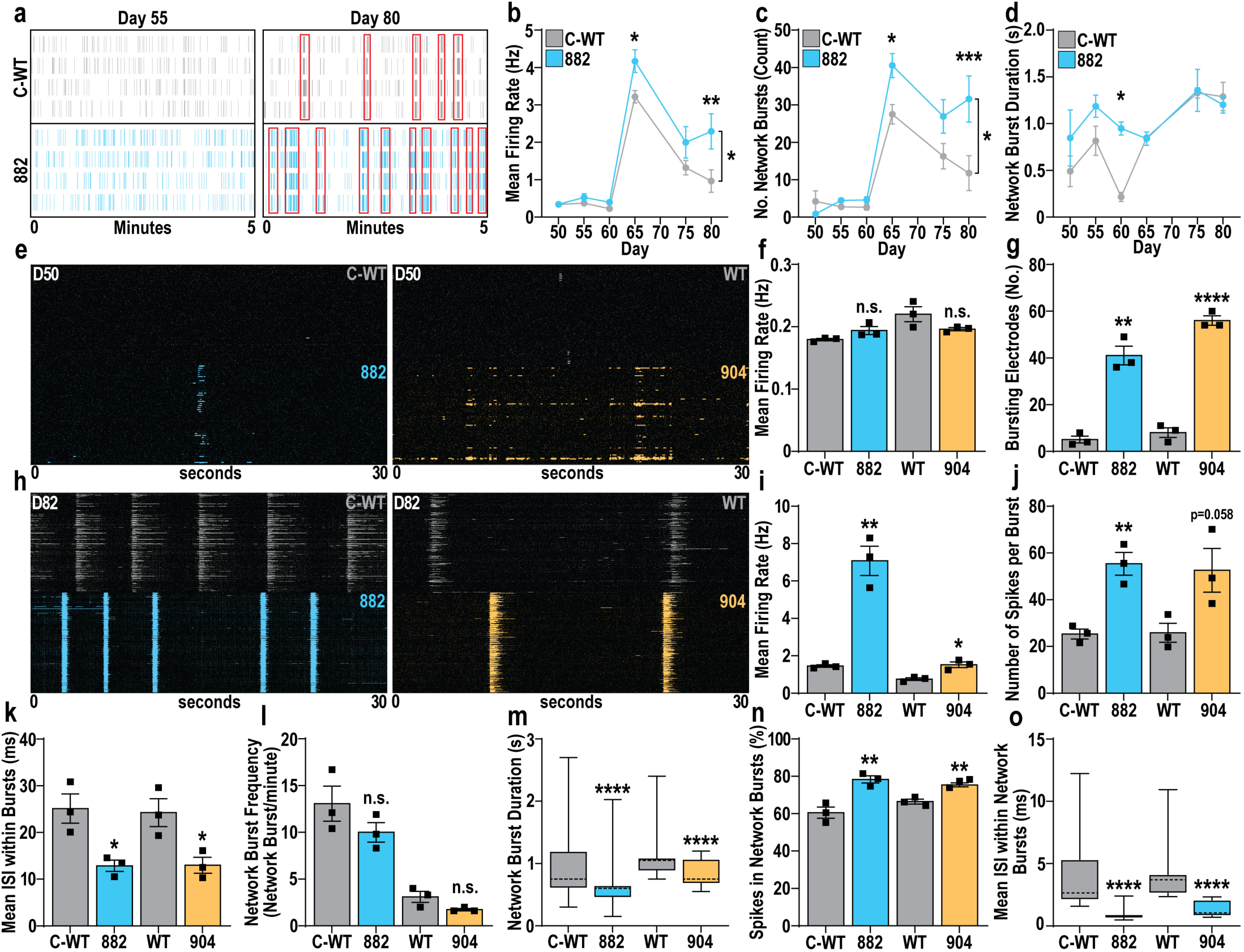
TBRS neuronal network dysfunction is driven by GABAergic neuron hyperactivity. **a,** Representative raster plots of LD-MEA activity in C-WT or 882 GABAergic/glutamatergic neuron co-cultures from early (D55, left) to late (D80, right) time points, with synchronized activity indicated by red boxes. **b-d,** Characteristics of C-WT and 882 neuronal networks, highlighting increased **(b)** firing rate (F_16_=6.579, pValue<0.05,n=4) and **(c)** frequency of network bursting (F_1_._6_=12.18,pValue<0.05,n=4) in 882 neuronal networks with no substantial changes in **(d)** network burst duration. **e-g,** Representative raster plots of HD-MEA activity **(e)** assessing the contribution of TBRS GABAergic neurons to neuronal network dysfunction after 50 days of differentiation, highlighting effects of TBRS GABAergic neurons on **(f)** firing rate and **(g)** appearance of neuronal bursting activity. **h-o,** HD-MEA activity assessing the contribution of TBRS GABAergic neurons to neuronal network dysfunction after 82 days of differentiation. **h-i,** Representative raster plots and quantification of changes in firing rate in cultures containing TBRS GABAergic neurons. **j-k,** Characterization of neuronal bursting parameters altered by TBRS GABAergic neurons, including U) the number of spikes per burst and the **(k)** inter-spike-interval (ISi) between spikes within neuronal bursts. **1-o,** Characterization of neuronal network parameters altered in the presence of TBRS GABAergic neurons, including network burst (I) frequency and **(m)** duration alongside the **(n)** proportion of activity within network bursts and the **(o)** ISi of spikes within network bursts. Data was analyzed by: two-way ANOVA, with posthoc Tukey’s multiple comparison testing **(b-d),**Student’s t-test **(f,g,i,j,k,l,n)** or Mann-Whitney test **(m,o),** versus isogenic controls.Data is represented as mean+/-SEM **(b-d,f,g,i,j,k,l,n)** with data points representing individual biological replicates **(f,g,i,j,k,l,n)** or as box-whisker plots with means indicated with dotted lines **(m,o).** For all experiments n=3 biological replicates; *pValue<0.05; **pValue<0.01; ***pValue<0.001; ****pValue<0.0001; n.s.=not significant.

Given the importance of GABAergic neurons in synchronizing neuronal activity (Makinen et al. 2018; Kirmse and Zhang 2022), we further investigated changes in neuronal bursting and neuronal network activity in these D82 co-cultures. This analysis highlighted decreased AP height (Supplemental Fig. S15c-e) in bursting neurons and increased numbers of spikes per burst (Fig. 7j), in cultures containing TBRS GABAergic neurons. Furthermore, cultures with 882 GABAergic neurons showed both increased bursting frequency (Supplemental Fig. S15f) and proportion of spikes in bursts with decreased inter-spike interval (ISI) within bursts (Fig. 7k, Supplemental Fig. S15g). Finally, while we did not observe alterations in the frequency of neuronal network activity associated with either TBRS model (Fig. 7l), TBRS GABAergic neurons caused a decreased duration of network bursts (Fig. 7m), coupled with increased numbers and proportions of spikes within network bursts (Fig. 7n, Supplemental Fig. S15h). These alterations were accompanied by decreased ISI and increased ISI coefficient of variation (CoV) of spikes within network bursts (Fig. 7o, Supplemental Fig. S15i). Together, these data demonstrate that both 882 and 904 TBRS-associated GABAergic neuron dysfunction is sufficient to drive neuronal network hyper-synchrony.

## Discussion

In this work, we derived new human models of TBRS and used these to characterize TBRS-associated alterations of neurodevelopment and neuronal function, identifying GABAergic neuron development as selectively sensitive to pathogenic *DNMT3A* mutation. We identified critical functions for DNMT3A in restraining both proliferation and neurogenesis during GABAergic neurodevelopment. We demonstrated that disruption of DNA methylation in TBRS models causes premature expression of neuronal and synaptic genes during GABAergic neuron differentiation, driving GABAergic neurogenesis and, ultimately, neuronal hyperactivity. This GABAergic neuron hyperactivity is sufficient to disrupt neuronal network formation, causing neuronal network hyper-synchronicity. Finally, this work highlighted mechanistic links between TBRS and OGIDs caused by mutations in other epigenetic modifiers and in the PIK3/AKT/mTOR signaling axis. Together, these findings reveal new developmental and lineage-specific roles for DNMT3A likely to underlie TBRS pathogenesis.

Significantly, the differential extent of TBRS-associated LOF in our models correlated with phenotypic severity, reminiscent of findings from the 882 and 904 orthologous TBRS mouse models (Smith et al. 2021; Beard et al. 2023). However, murine TBRS studies to date have focused on non-CG methylation catalyzed by DNMT3A in post-mitotic neurons of adult animals (Christian et al. 2020; Beard et al. 2023), while we instead found that aberrant development of GABAergic neurons underlies neuronal network hyperactivity and hypersynchrony in human TBRS models. Similarly, DNMT3A homozygous knockout during human motor neuron (hMN) development altered hMN function, causing hyperactivity; however, this was accompanied by impaired neurogenesis and synapse production (Ziller et al. 2018), counter to our findings. Our findings instead demonstrate a role for DNMT3A in restraining cortical neuron maturation, defining distinct cell-type specific consequences of DNMT3A disruption by TBRS mutation. We also identified hyperproliferation specific to GABAergic neuron specification across TBRS models, which may contribute to the brain overgrowth in TBRS patients. Given the lack of brain overgrowth reported in murine models of TBRS (Christian et al. 2020; Beard et al. 2023), our work suggests that this may be a human specific phenomenon, consistent with findings for several other neurodevelopmental disorders (Ernst 2016; Marchetto et al. 2017; Wang et al. 2020; Connacher et al. 2022). Our results also support functional interplay between TBRS and PROS, with TBRS models showing increased PIK3/AKT/mTOR signaling, consistent with the finding that gain of function mutations in components of this pathway cause the related brain overgrowth disorder PROS (Mirzaa et al. 2016). While PROS-associated mutations remain understudied (Mirzaa et al. 2016; Dobyns and Mirzaa 2019) this signaling axis promotes excessive proliferation in the context of cancer (Samuels et al. 2005; Wang et al. 2017; Chen et al. 2022), such that pharmacological modifiers of PIK3/AKT/mTOR signaling are widely available. While our work highlighted the use of rapamycin in modulating hyperproliferation, prior studies have used other modulators of this signaling axis to rescue neuronal network abnormalities associated with TSC deficiency (Alsaqati et al. 2020), which likewise disrupts GABAergic neuron differentiation (Fu et al. 2012) and function (Bassetti et al. 2021). Therefore, if altered PIK3/AKT/mTOR signaling is a common hallmark across OGIDs, as our data and that of others (Dai et al. 2017) suggests, this may present an opportunity for identifying common mechanistic intervention paradigms to treat OGIDs originating from genetically distinct mutations.

Our findings also support mechanistic interplay between EZH2 and DNMT3A in restraining human GABAergic neuron differentiation, findings congruent with a functional relationship between EZH2 and DNMT3A reported in murine models (Li et al. 2022), thus suggesting a shared mechanism between the OGIDs TBRS and Weaver syndrome (caused by *EZH2* mutations) (Tatton-Brown et al. 2013; Cohen et al. 2016; Tatton-Brown et al. 2017). Recent work has demonstrated a role for EZH2 in restraining glutamatergic neuron maturation (Ciceri et al. 2024), suggesting a critical role for other OGID-associated genes in regulating neuronal maturation. Similarly, neuronal maturation is disrupted in human models of pathogenic *MECP2* mutation modeling Rett Syndrome (Landucci et al. 2018), which disrupts MECP2 read out of DNMT3A-deposited DNA methylation (Clemens et al. 2020; Sandweiss et al. 2020; Pantier et al. 2024). Furthermore, restoring MECP2 expression specifically in GABAergic neurons in Rett mouse models rescued multiple Rett syndrome-associated phenotypes (Ure et al. 2016). Alongside our results, these findings confirm a selective sensitivity of GABAergic neurons to neurodevelopmental disorder-associated perturbations that disrupt epigenetic repression of neuronal maturation.

Our HD-MEA results further demonstrate that GABAergic neuron dysfunction is sufficient to imbalance excitatory and inhibitory signaling in TBRS, while such E/I imbalance is a common mechanism suggested to underlie pathogenesis in neurodevelopmental disorders (Gatto and Broadie 2010; Uzunova et al. 2016; Canitano and Pallagrosi 2017; Markicevic et al. 2020; Pietropaolo and Provenzano 2022). Although MEA assessments of neurodevelopmental disorder models are becoming more prevalent, conclusions from LD-MEA studies are constrained by a limited quantity of data, often resulting in inconsistent findings both within and across neurodevelopmental disorder studies (Nageshappa et al. 2016; Marchetto et al. 2017; Amatya et al. 2019; Deneault et al. 2019a; Deneault et al. 2019b; Frega et al. 2019; Alsaqati et al. 2020; Graef et al. 2020; Chapman et al. 2022; DeRosa et al. 2022; Rylaarsdam et al. 2024). Therefore, while LD-MEA assessments in cellular models of Rett Syndrome revealed some phenotypic similarities with our TBRS models (Mok et al. 2022), our study highlights the importance of using HD-MEA to resolve inconsistencies driven by limited data in LD-MEA experiments. Ultimately, the paradigm we present here using HD-MEA provides a high-quality workflow for assessing the role of altered GABAergic neuron function in disrupting neuronal network development, which can be applied across future studies of OGIDs and other NDDs.

In summary, we found that TBRS-associated GABAergic neuron dysfunction is sufficient to disrupt the development of neuronal networks and is likely to contribute to the etiology of the ASD and ID common among TBRS patients. Furthermore, our findings of molecular and phenotypic relationships between different OGIDs highlight convergent mechanisms that may underlie their shared clinical presentation. Together, these results suggest that future efforts to develop interventions for TBRS may have broader applicability across OGIDs, while providing significant information on sensitive cell types and developmental periods that will facilitate the development of such interventions.

## Methods

### hPSC Model Generation and Culture

Work with hPSCs was performed in accordance with the Washington University Embryonic Stem Cell Research Oversight Committee (ESCRO) under protocol #12-002. DNMT3A R882H iPSCs were reprogrammed from a male patient carrying heterozygous p.R882H (c.2922G>A) mutation and corrected using CRISPR-Cas9 technology. Whole genome sequencing was performed on both the R882H and subsequently corrected control (C-WT) iPSC lines, with summary data in Supplementary Data 1 and full data available from dbGaP (PHS000159). Production of the three primordial germ layers from both R882H and C-WT iPSCs was confirmed by assessing teratoma formation upon iPSC injection into the mouse fat pad.

Del1 and Del2 iPSC models were similarly generated from a male patient with a heterozygous 135 kb deletion containing the entire *DNMT3A* gene. The associated controls (Con1/2) were purchased from FUJIFILM Cellular Dynamics, Inc. (CW20110 and CW20098) as sex matched unrelated controls with no history of neurodevelopmental disorders including autism spectrum disorder and epilepsy. The heterozygous P904L (c.2711C>T) mutation was modelled by its heterozygous introduction into H1 human embryonic stem cells (WT) using CRISPR-Cas9, with the same process generating a model homozygous for small (<10bp) deletions in both alleles of the DNMT3A gene centered on position c.2711 in the cDNA (KO).

DNMT3A CRISPRi lines were created by transducing H1 hESCs with a single vector (Lenti-(BB)-EF1a-KRAB-dCas9-P2A-BlastR) carrying expression cassettes for both dCas9-KRAB and each gRNA. gRNAs (Supplementary Table 1) for DNMT3A from the Dolcetto library (Sanson et al. 2018) were cloned into Lenti-(BB)-EF1a-KRAB-dCas9-P2A-BlastR. Cell lines used to produce induced glutamatergic-like neurons (iGluts) were generated by transducing cell lines with lentivirus carrying pLVX-UbC-rtTA- Ngn2:2A:EGFP. For full details on lentivirus production and stable line generation see Supplementary material.

All hPSC models were maintained under feeder-free conditions on vitronectin in StemFlex Medium and cultured in an incubator with 5% CO2 at 37°C throughout all experiments. Experiments were carried out between passages 20-50 for all hPSC lines and lines periodically tested as negative for mycoplasma contamination, were confirmed karyotypically normal (Supplementary Fig. S17a) and validated for pluripotency markers (OCT4, SOX2, Supplementary Fig. S17b-c).

### Modeling cortical GABAergic and glutamatergic neuron development

Modeling of cortical GABAergic neuron development was carried out as previously described (Chapman et al. 2024) by specifying hPSCs as medial ganglionic eminence (MGE)-like progenitors with a ventral telencephalic progenitor (V-NPCs) character and differentiating V-NPCs into immature GABAergic cortical interneurons (V-INs). Cells were collected on day 15 of differentiation as V-NPCs and on day 35 of differentiation as V-INs. Further maturation of V-INs was performed for functional assessments as described below. For full details, see Supplementary Material.

Modeling of cortical glutamatergic neuron development was carried out as previously described (Chapman et al. 2022) with minor modifications by specifying hPSCs as subventricular zone-like progenitors with a dorsal telencephalic progenitor character (D-NPCs) and differentiating these D-NPCs into immature glutamatergic neurons (D-INs). Cells were collected on day 20 of differentiation as D-NPCs and on day 40 of differentiation as D-INs. Further maturation of D-INs was performed for functional assessments as described below. For full details, see Supplementary Material.

Induced glutamatergic-like neurons were generated as previously described (Schafer et al. 2019) by inducing overexpression of Neurogenin-2 (NGN2). Cells were considered iGluts after 14 days of differentiation. For full details see Supplementary Material.

Organoids (Orgs) with a dorsal (D) or ventral (V) telencephalic character were generated by embedding patterned neurospheres, generated during D-NPC or V-NPC specification in Matrigel®. D- and V-Orgs were then maintained on an orbital shaker for a total of 30 days of differentiation; for full details, see Supplementary Material. Organoids were collected and fixed in 4% Paraformaldehyde overnight at 4℃ before being incubated in 30% (v/v) sucrose solution overnight at 4℃. Organoids were embedded in a 1:1 ratio of O.C.T compound and 30% sucrose before being sectioned on a cryostat at 8 µm.

### Cellular Phenotyping

Changes in DNMT3A expression across TBRS models was assessed by western blot using protein isolated from pluripotent stem cells with full blots shown in Supplemental Fig. S16. Quantifications of neurosphere outgrowth were performed in D-NPCs and V-NPCs after 12 days of differentiation (D12) and changes in proliferation and lineage marker expression were assessed by immunocytochemistry 4 days after D-NPC or V-NPC specification was complete. Analogous quantifications of organoid size, proliferation, and lineage marker expression were made in D-Orgs and V-Orgs after 30 days of differentiation. RT-qPCR to assess changes in the expression of specific genes was also performed from D-NPC and V-NPC samples isolated after specification was complete.

Changes in PIK3/AKT/mTOR signaling were assessed by western blotting using V-NPC samples, and rapamycin (5 nM) was applied to V-NPCs for 4 days after V-NPC specification was complete to assess changes in proliferation upon mTOR inhibition. Similarly, the effects of EZH2 inhibition were assessed by RT-qPCR in D-NPCs treated with EZH2i (GSK343 at 4 µM) versus DMSO or siRNAs targeting EZH2 versus siRNA for GFP for 4 days after specification was complete.

TBRS-associated alterations to neuronal differentiation were assessed by immunocytochemistry for markers of neuronal differentiation in D-Orgs and V-Orgs at D30, with similar quantifications made in V-INs at D40. Neuronal morphology was assessed in V-INs at D30, and synapse production of V-INs was assessed at D50 after V-INs were grown in co-culture with rat astrocytes for 20 days. For further information on the methods described above, see Supplementary Material.

### Sequencing

RNA sequencing (RNA-seq) and whole genome bisulfite sequencing (WGBS) was performed as previously described (Hamagami et al. 2023; Chapman et al. 2024) on 882, 904, C-WT, and WT V-NPCs at D15, with similar analysis performed in 882 and C-WT D-NPCs at D20. CUT&Tag for histone H3 lysine 27 tri-methylation was performed as previously described (Chapman et al. 2024) on 882, 904, C-WT and WT V-NPCs at D15. RNA-seq and WGBS were also performed in 882, 904, C-WT and WT D-and V-INs at D40 and D35 respectively with similar analysis performed in 882, 904, C-WT and WT iGluts at D14. While the protocols used are able to generate populations with >85-90% purity for lineage-specific marker expression, pathogenic mutations may disrupt the specification and differentiation timing of these cells, therefore, all -omics data was generated from these mixed cell populations. For full details on CUT&Tag, RNA-seq and WGBS methodologies see Supplementary Material.

### Functional Phenotyping

For electrophysiology, D28 D-INs and D23 V-INs were plated on rat cortical astrocytes and matured until D60, before electrophysiological measurements were generated as described previously (Meganathan et al. 2017). Results were analyzed using non-parametric rank sum tests comparing TBRS models to matched isogenic controls with raw results presented in Supplemental Data.

All low-density multi-electrode array (MEA) experiments were performed using CytoView MEA 48 well plates using co-cultures of matured D-INs and V-INs (at a 70:30 ratio). Electrophysiological activity was recorded every 5 days using hardware (Maestro Edge) and software (AxIS 1.5.2) from Axion Biosystems, with analysis performed using the built-in Axion neural metric analysis tool.

All high-density MEA experiments were performed using CorePlate™ 6W 38/60 MEAs using co-cultures of V-INs and iGluts (at a 30:70 ratio) alongside rat astrocytes. Electrophysiological activity was recorded on D50 and D82 of V-IN differentiation, using the HyperCAM Alpha system (3Brain) and accompanying BrainWave5 software, with primary data analysis performed using BrainWave5 software. For full details on electrophysiological and MEA characterization of TBRS models, see Supplementary Material.

### Quantification and Statistical Analysis

Where appropriate, statistical analysis was carried out using GraphPad Prism version 9 (GraphPad Software; La Jolla, CA, USA, available from www.graphpad.com) and/or RStudio version 3.5.1 (RStudio: Integrated development environment for R; Boston, MA, USA. Available from www.rstudio.org). All technical replicates were averaged before statistical analysis, and statistical tests used for each data analysis are detailed in the figure legends or in detailed methods section for specific analysis paradigms. A minimum of 3 independent differentiations were used for each time point or biological condition, with the number of differentiations used for each sample listed in figure legends as n. The results in figures are presented as group mean +/- standard error (SE), indicating each biological replicate used for the analysis unless otherwise specified in figure legends. Statistical significance is indicated as follows: n.s., not significant; ∗, p < 0.05; ∗∗, p < 0.01; ∗∗∗, p < 0.001; ∗∗∗∗, p < 0.0001, unless otherwise specified in figure legends.

## Datasets

Raw and processed data was deposited into the Gene Expression Omnibus as accession numbers GSE294191 (DNA methylation), GSE294189 (RNA sequencing) and GSE294186 (H3K27me3 CUT&Tag).

## Acknowledgements

This work was supported by NIH Grants R01NS114551, R01MH124808, R01HD110556, and U0-HG007530 (NIH Common Fund/NINDS Undiagnosed Disease Network pilot gene study subaward) to KLK, and by NIH P50HD103525 to Joseph Dougherty and Christina Gurnett (KLK is project PI for the Human Cellular Models Unit of the Washington University Intellectual and Developmental Disabilities Research Center, with project effort funded by this grant). Foundations supporting this work included the Engelhart Family Foundation, Simons Foundation, M-CM Network, and pilot awards from the WU Hope Center, Center of Regenerative Medicine, and Institute for Clinical and Translational Sciences to KLK

We thank the Genome Technology Access Center at the McDonnell Genome Institute at Washington University School of Medicine for help with genomics services. The Center is partially supported by NCI Cancer Center Support Grant #P30 CA91842 to the Siteman Cancer Center from the National Center for Research Resources (NCRR), a component of the National Institutes of Health (NIH) and NIH Roadmap for Medical Research. This publication is solely the responsibility of the authors and does not necessarily represent the official view of NCRR or NIH. We also thank the Genome Engineering & Stem Cell Center (GESC@MGI) at Washington University in St. Louis for cell line engineering services. The DNMT3A R882H iPSC line and Del1/2 iPSC lines used in this study were developed and characterized in the laboratory of Timothy J. Ley at Washington University, by Daniel George and Dr. Christopher A. Miller, supported by NIH CA197161.

## Author Contributions

G.C. and J.J.D. performed differentiations and MEA assessments and analyzed all results. G.C., J.R.E., and T.E.L. performed computational and bioinformatics analysis. J.E.H and J.J.D. performed electrophysiological assessments and associated analysis. G.C. and S.R.C. performed qPCR and associated analysis. G.C., F.B., R.P., S.M., and H.J. performed immunocytochemistry assessments, Y.R. prepared samples for bi-sulfite sequencing, and G.C. and F.B. performed western blotting analysis. H.W.G. and K.L.K. supervised the work. G.C. wrote the manuscript, and all the authors contributed to reviewing and editing the manuscript.

## References

Alsaqati M, Heine VM, Harwood AJ. 2020. Pharmacological intervention to restore connectivity deficits of neuronal networks derived from ASD patient iPSC with a TSC2 mutation. Mol Autism 11: 80.

Amatya DN, Linker SB, Mendes APD, Santos R, Erikson G, Shokhirev MN, Zhou Y, Sharpee T, Gage FH, Marchetto MC et al. 2019. Dynamical Electrical Complexity Is Reduced during Neuronal Differentiation in Autism Spectrum Disorder. Stem Cell Reports 13: 474–484.

Atterton C, Trew I, Cale JM, Aung-Htut MT, Grens K, Kiernan J, Delagrammatikas CG, Piper M. 2025. Overgrowth-intellectual disability disorders: progress in biology, patient advocacy and innovative therapies. Dis Model Mech 18.

Bassetti D, Luhmann HJ, Kirischuk S. 2021. Effects of Mutations in TSC Genes on Neurodevelopment and Synaptic Transmission. Int J Mol Sci 22.

Beard DC, Zhang X, Wu DY, Martin JR, Erickson A, Boua JV, Hamagami N, Swift RG, McCullough KB, Ge X et al. 2023. Distinct disease mutations in DNMT3A result in a spectrum of behavioral, epigenetic, and transcriptional deficits. Cell Rep 42: 113411.

Canitano R, Pallagrosi M. 2017. Autism Spectrum Disorders and Schizophrenia Spectrum Disorders: Excitation/Inhibition Imbalance and Developmental Trajectories. Front Psychiatry 8: 69.

Chapman G, Alsaqati M, Lunn S, Singh T, Linden SC, Linden DEJ, van den Bree MBM, Ziller M, Owen MJ, Hall J et al. 2022. Using induced pluripotent stem cells to investigate human neuronal phenotypes in 1q21.1 deletion and duplication syndrome. Mol Psychiatry 27: 819–830.

Chapman G, Determan J, Jetter H, Kaushik K, Prakasam R, Kroll KL. 2024. Defining cis-regulatory elements and transcription factors that control human cortical interneuron development. iScience 27: 109967.

Chen W, Dai G, Qian Y, Wen L, He X, Liu H, Gao Y, Tang X, Dong B. 2022. PIK3CA mutation affects the proliferation of colorectal cancer cells through the PI3K-MEK/PDK1-GPT2 pathway. Oncol Rep 47.

Christian DL, Wu DY, Martin JR, Moore JR, Liu YR, Clemens AW, Nettles SA, Kirkland NM, Papouin T, Hill CA et al. 2020. DNMT3A Haploinsufficiency Results in Behavioral Deficits and Global Epigenomic Dysregulation Shared across Neurodevelopmental Disorders. Cell Rep 33: 108416.

Ciceri G, Baggiolini A, Cho HS, Kshirsagar M, Benito-Kwiecinski S, Walsh RM, Aromolaran KA, Gonzalez-Hernandez AJ, Munguba H, Koo SY et al. 2024. An epigenetic barrier sets the timing of human neuronal maturation. Nature 626: 881–890.

Clemens AW, Wu DY, Moore JR, Christian DL, Zhao G, Gabel HW. 2020. MeCP2 Represses Enhancers through Chromosome Topology-Associated DNA Methylation. Mol Cell 77: 279–293 e278.

Cohen AS, Yap DB, Lewis ME, Chijiwa C, Ramos-Arroyo MA, Tkachenko N, Milano V, Fradin M, McKinnon ML, Townsend KN et al. 2016. Weaver Syndrome-Associated EZH2 Protein Variants Show Impaired Histone Methyltransferase Function In Vitro. Hum Mutat 37: 301–307.

Connacher R, Williams M, Prem S, Yeung PL, Matteson P, Mehta M, Markov A, Peng C, Zhou X, McDermott CR et al. 2022. Autism NPCs from both idiopathic and CNV 16p11.2 deletion patients exhibit dysregulation of proliferation and mitogenic responses. Stem Cell Reports 17: 1786.

Dai YJ, Wang YY, Huang JY, Xia L, Shi XD, Xu J, Lu J, Su XB, Yang Y, Zhang WN et al. 2017. Conditional knockin of Dnmt3a R878H initiates acute myeloid leukemia with mTOR pathway involvement. Proc Natl Acad Sci U S A 114: 5237–5242.

Deneault E, Faheem M, White SH, Rodrigues DC, Sun S, Wei W, Piekna A, Thompson T, Howe JL, Chalil L et al. 2019a. CNTN5(-)(/+)or EHMT2(-)(/+)human iPSC-derived neurons from individuals with autism develop hyperactive neuronal networks. Elife 8.

Deneault E, White SH, Rodrigues DC, Ross PJ, Faheem M, Zaslavsky K, Wang Z, Alexandrova R, Pellecchia G, Wei W et al. 2019b. Complete Disruption of Autism-Susceptibility Genes by Gene Editing Predominantly Reduces Functional Connectivity of Isogenic Human Neurons. Stem Cell Reports 12: 427–429.

DeRosa BA, Hokayem JE, Artimovich E, Garcia-Serje C, Phillips AW, Van Booven D, Nestor JE, Wang L, Cuccaro ML, Vance JM et al. 2022. Author Correction: Convergent Pathways in Idiopathic Autism Revealed by Time Course Transcriptomic Analysis of Patient-Derived Neurons. Sci Rep 12: 3445.

Dobyns WB, Mirzaa GM. 2019. Megalencephaly syndromes associated with mutations of core components of the PI3K-AKT-MTOR pathway: PIK3CA, PIK3R2, AKT3, and MTOR. Am J Med Genet C Semin Med Genet 181: 582–590.

Ernst C. 2016. Proliferation and Differentiation Deficits are a Major Convergence Point for Neurodevelopmental Disorders. Trends Neurosci 39: 290–299.

Feng J, Chang H, Li E, Fan G. 2005. Dynamic expression of de novo DNA methyltransferases Dnmt3a and Dnmt3b in the central nervous system. J Neurosci Res 79: 734–746.

Frega M, Linda K, Keller JM, Gumus-Akay G, Mossink B, van Rhijn JR, Negwer M, Klein Gunnewiek T, Foreman K, Kompier N et al. 2019. Neuronal network dysfunction in a model for Kleefstra syndrome mediated by enhanced NMDAR signaling. Nat Commun 10: 4928.

Fu C, Cawthon B, Clinkscales W, Bruce A, Winzenburger P, Ess KC. 2012. GABAergic interneuron development and function is modulated by the Tsc1 gene. Cereb Cortex 22: 2111–2119.

Gatto CL, Broadie K. 2010. Genetic controls balancing excitatory and inhibitory synaptogenesis in neurodevelopmental disorder models. Front Synaptic Neurosci 2: 4.

Gibson WT, Hood RL, Zhan SH, Bulman DE, Fejes AP, Moore R, Mungall AJ, Eydoux P, Babul-Hirji R, An J et al. 2012. Mutations in EZH2 cause Weaver syndrome. Am J Hum Genet 90: 110–118.

Graef JD, Wu H, Ng C, Sun C, Villegas V, Qadir D, Jesseman K, Warren ST, Jaenisch R, Cacace A et al. 2020. Partial FMRP expression is sufficient to normalize neuronal hyperactivity in Fragile X neurons. Eur J Neurosci 51: 2143–2157.

Hamagami N, Wu DY, Clemens AW, Nettles SA, Li A, Gabel HW. 2023. NSD1 deposits histone H3 lysine 36 dimethylation to pattern non-CG DNA methylation in neurons. Molecular Cell 83: 1412–1428.e1417.

Kang HJ, Kawasawa YI, Cheng F, Zhu Y, Xu X, Li M, Sousa AM, Pletikos M, Meyer KA, Sedmak G et al. 2011. Spatio-temporal transcriptome of the human brain. Nature 478: 483–489.

Kirmse K, Zhang C. 2022. Principles of GABAergic signaling in developing cortical network dynamics. Cell Rep 38: 110568.

Landucci E, Brindisi M, Bianciardi L, Catania LM, Daga S, Croci S, Frullanti E, Fallerini C, Butini S, Brogi S et al. 2018. iPSC-derived neurons profiling reveals GABAergic circuit disruption and acetylated alpha-tubulin defect which improves after iHDAC6 treatment in Rett syndrome. Exp Cell Res 368: 225–235.

Lane C, Tatton-Brown K, Freeth M. 2020. Tatton-Brown-Rahman syndrome: cognitive and behavioural phenotypes. Dev Med Child Neurol 62: 993–998.

Li J, Pinto-Duarte A, Zander M, Cuoco MS, Lai CY, Osteen J, Fang L, Luo C, Lucero JD, Gomez-Castanon R et al. 2022. Dnmt3a knockout in excitatory neurons impairs postnatal synapse maturation and increases the repressive histone modification H3K27me3. Elife 11.

Makinen ME, Yla-Outinen L, Narkilahti S. 2018. GABA and Gap Junctions in the Development of Synchronized Activity in Human Pluripotent Stem Cell-Derived Neural Networks. Front Cell Neurosci 12: 56.

Marchetto MC, Belinson H, Tian Y, Freitas BC, Fu C, Vadodaria K, Beltrao-Braga P, Trujillo CA, Mendes APD, Padmanabhan K et al. 2017. Altered proliferation and networks in neural cells derived from idiopathic autistic individuals. Mol Psychiatry 22: 820–835.

Markicevic M, Fulcher BD, Lewis C, Helmchen F, Rudin M, Zerbi V, Wenderoth N. 2020. Cortical Excitation:Inhibition Imbalance Causes Abnormal Brain Network Dynamics as Observed in Neurodevelopmental Disorders. Cereb Cortex 30: 4922–4937.

Meganathan K, Lewis EMA, Gontarz P, Liu S, Stanley EG, Elefanty AG, Huettner JE, Zhang B, Kroll KL. 2017. Regulatory networks specifying cortical interneurons from human embryonic stem cells reveal roles for CHD2 in interneuron development. Proc Natl Acad Sci U S A 114: E11180–E11189.

Mirzaa G, Timms AE, Conti V, Boyle EA, Girisha KM, Martin B, Kircher M, Olds C, Juusola J, Collins S et al. 2016. PIK3CA-associated developmental disorders exhibit distinct classes of mutations with variable expression and tissue distribution. JCI Insight 1.

Mok RSF, Zhang W, Sheikh TI, Pradeepan K, Fernandes IR, DeJong LC, Benigno G, Hildebrandt MR, Mufteev M, Rodrigues DC et al. 2022. Wide spectrum of neuronal and network phenotypes in human stem cell-derived excitatory neurons with Rett syndrome-associated MECP2 mutations. Transl Psychiatry 12: 450.

Nageshappa S, Carromeu C, Trujillo CA, Mesci P, Espuny-Camacho I, Pasciuto E, Vanderhaeghen P, Verfaillie CM, Raitano S, Kumar A et al. 2016. Altered neuronal network and rescue in a human MECP2 duplication model. Mol Psychiatry 21: 178–188.

Nguyen TV, Yao S, Wang Y, Rolfe A, Selvaraj A, Darman R, Ke J, Warmuth M, Smith PG, Larsen NA et al. 2019. The R882H DNMT3A hot spot mutation stabilizes the formation of large DNMT3A oligomers with low DNA methyltransferase activity. J Biol Chem 294: 16966–16977.

Ostrowski PJ, Tatton-Brown K. 1993. Tatton-Brown-Rahman Syndrome. in GeneReviews((R)) (eds. MP Adam, J Feldman, GM Mirzaa, RA Pagon, SE Wallace, LJH Bean, KW Gripp, A Amemiya), Seattle (WA).

Pantier R, Brown M, Han S, Paton K, Meek S, Montavon T, Shukeir N, McHugh T, Kelly DA, Hochepied T et al. 2024. MeCP2 binds to methylated DNA independently of phase separation and heterochromatin organisation. Nat Commun 15: 3880.

Pietropaolo S, Provenzano G. 2022. Editorial: Targeting Excitation-Inhibition Imbalance in Neurodevelopmental and Autism Spectrum Disorders. Front Neurosci 16: 968115.

Romanyuk N, Sintakova K, Arzhanov I, Horak M, Gandhi C, Jhanwar-Uniyal M, Jendelova P. 2024. mTOR pathway inhibition alters proliferation as well as differentiation of neural stem cells. Front Cell Neurosci 18: 1298182.

Rylaarsdam L, Rakotomamonjy J, Pope E, Guemez-Gamboa A. 2024. iPSC-derived models of PACS1 syndrome reveal transcriptional and functional deficits in neuron activity. Nat Commun 15: 827.

Samuels Y, Diaz LA, Jr., Schmidt-Kittler O, Cummins JM, Delong L, Cheong I, Rago C, Huso DL, Lengauer C, Kinzler KW et al. 2005. Mutant PIK3CA promotes cell growth and invasion of human cancer cells. Cancer Cell 7: 561–573.

Sandweiss AJ, Brandt VL, Zoghbi HY. 2020. Advances in understanding of Rett syndrome and MECP2 duplication syndrome: prospects for future therapies. Lancet Neurol 19: 689–698.

Sanson KR, Hanna RE, Hegde M, Donovan KF, Strand C, Sullender ME, Vaimberg EW, Goodale A, Root DE, Piccioni F et al. 2018. Optimized libraries for CRISPR-Cas9 genetic screens with multiple modalities. Nat Commun 9: 5416.

Schafer ST, Paquola ACM, Stern S, Gosselin D, Ku M, Pena M, Kuret TJM, Liyanage M, Mansour AA, Jaeger BN et al. 2019. Pathological priming causes developmental gene network heterochronicity in autistic subject-derived neurons. Nat Neurosci 22: 243–255.

Smith AM, LaValle TA, Shinawi M, Ramakrishnan SM, Abel HJ, Hill CA, Kirkland NM, Rettig MP, Helton NM, Heath SE et al. 2021. Functional and epigenetic phenotypes of humans and mice with DNMT3A Overgrowth Syndrome. Nat Commun 12: 4549.

Tatton-Brown K, Loveday C, Yost S, Clarke M, Ramsay E, Zachariou A, Elliott A, Wylie H, Ardissone A, Rittinger O et al. 2017. Mutations in Epigenetic Regulation Genes Are a Major Cause of Overgrowth with Intellectual Disability. Am J Hum Genet 100: 725–736.

Tatton-Brown K, Murray A, Hanks S, Douglas J, Armstrong R, Banka S, Bird LM, Clericuzio CL, Cormier-Daire V, Cushing T et al. 2013. Weaver syndrome and EZH2 mutations: Clarifying the clinical phenotype. Am J Med Genet A 161A: 2972–2980.

Tatton-Brown K, Zachariou A, Loveday C, Renwick A, Mahamdallie S, Aksglaede L, Baralle D, Barge-Schaapveld D, Blyth M, Bouma M et al. 2018. The Tatton-Brown-Rahman Syndrome: A clinical study of 55 individuals with de novo constitutive DNMT3A variants. Wellcome Open Res 3: 46.

Thomas H, Alix T, Renard E, Renaud M, Wourms J, Zuily S, Leheup B, Genevieve D, Dreumont N, Schmitt E et al. 2024. Expanding the genetic and clinical spectrum of Tatton-Brown-Rahman syndrome in a series of 24 French patients. J Med Genet.

Ure K, Lu H, Wang W, Ito-Ishida A, Wu Z, He LJ, Sztainberg Y, Chen W, Tang J, Zoghbi HY. 2016. Restoration of Mecp2 expression in GABAergic neurons is sufficient to rescue multiple disease features in a mouse model of Rett syndrome. Elife 5.

Uzunova G, Pallanti S, Hollander E. 2016. Excitatory/inhibitory imbalance in autism spectrum disorders: Implications for interventions and therapeutics. World J Biol Psychiatry 17: 174–186.

Wang L, Huang D, Jiang Z, Luo Y, Norris C, Zhang M, Tian X, Tang Y. 2017. Akt3 is responsible for the survival and proliferation of embryonic stem cells. Biol Open 6: 850–861.

Wang M, Wei PC, Lim CK, Gallina IS, Marshall S, Marchetto MC, Alt FW, Gage FH. 2020. Increased Neural Progenitor Proliferation in a hiPSC Model of Autism Induces Replication Stress-Associated Genome Instability. Cell Stem Cell 26: 221–233 e226.

Watanabe D, Uchiyama K, Hanaoka K. 2006. Transition of mouse de novo methyltransferases expression from Dnmt3b to Dnmt3a during neural progenitor cell development. Neuroscience 142: 727–737.

Wu Z, Huang K, Yu J, Le T, Namihira M, Liu Y, Zhang J, Xue Z, Cheng L, Fan G. 2012. Dnmt3a regulates both proliferation and differentiation of mouse neural stem cells. J Neurosci Res 90: 1883–1891.

Ziller MJ, Ortega JA, Quinlan KA, Santos DP, Gu H, Martin EJ, Galonska C, Pop R, Maidl S, Di Pardo A et al. 2018. Dissecting the Functional Consequences of De Novo DNA Methylation Dynamics in Human Motor Neuron Differentiation and Physiology. Cell Stem Cell 22: 559–574 e559.

