## Supplementary Materials, Figures and Tables for "Tatton-Brown-Rahman Syndrome-associated *DNMT3A* mutations de-repress cortical interneuron differentiation to disrupt neuronal network function"

This PDF file includes:


1. Supplementary Methods
2. Supplementary Tables
3. Supplementary Figures
4. References

**Supplementary Methods**

**Lentivirus production and stable line generation**

Lentivirus was produced in Lenti-X-293T cells (Takara Bio), by transfecting the cells with envelope (pMD2.G), packaging (psPAX2), and transfer plasmid (Lenti-(BB)-EF1a-KRAB-dCas9-P2A-BlastR, pLVX-UbC-rtTA-Ngn2:2A:EGFP or pHAGE-EZH2) at a ratio of 2:3:5 using Trans-IT Lenti transfection reagent (Mirus). pMD2.G and psPAX2 were gifts from Didier Trono (Addgene plasmids #12259 and #12260, respectively), Lenti-(BB)-EF1a-KRAB-dCas9-P2A-BlastR was a gift from Jorge Ferrer (Addgene plasmid # 118154), pLVX-UbC-rtTA-Ngn2:2A:EGFP was a gift from Fred Gage (Addgene plasmid # 127288) and pHAGE-EZH2 was a gift from Gordon Mills & Kenneth Scott (Addgene plasmid # 116738). Lenti-X-293T cells were grown in HEK media containing DMEM, high glucose, and no glutamine (Gibco), and supplemented with 10% FBS (Gibco), Glutamax, β-Mercaptoethanol, Non-Essential Amino Acids and Penicillin-Streptomycin Solution, Sodium Pyruvate (Life Technologies) and HEPES buffer (Life Technologies). Lentivirus was concentrated using Lenti-X-Concentrator (Takara Bio) per the manufacturer's instructions, re-suspended in Opti-MEM media (Gibco), and stored at -80°C.

Transduction of human pluripotent stem cell models was aided by the addition of 5 µg/mL polybrene (Sigma-Aldrich) immediately before the addition of lentivirus. Cells recovered for 48 hours before selection (500 ng/mL puromycin or 1 µg/mL blasticidin) was applied. Once cells were selected, they were maintained under selection for the duration of subsequent experiments for CRISPRi models or until D6 of iGlut differentiation.

**Ventral telencephalic progenitor specification and cortical GABAergic neuron differentiation**

Embryoid bodies (EBs) were produced from hPSCs using AggreWell™800 Microwell Culture Plates (STEMCELL Technologies) and AggreWell™ Embryoid Body Formation Medium (STEMCELL Technologies) and maintained for 3 days (day -3 to 0). Differentiation was then initiated at (day (D) 0) by switching to ventral telencephalic neuroectoderm specification media (VSM). VSM contained Neurobasal-A (Life Technologies), B-27 supplement (without Vitamin A; Life Technologies), Glutamax (Life Technologies), β-Mercaptoethanol (Life Technologies), Non-Essential Amino Acids (Life Technologies), Penicillin-Streptomycin Solution (Life Technologies), 10 µM SB-431542 (Selleckchem), 100 nM LDN-193189 (Tocris Biosciences), 100 nM Smoothened Agonist (SAG; Selleckchem), and 2 µM XAV-939 (Selleckchem). For the first 2 days of differentiation, media also contained 10 µM Y-27632 (Selleckchem). EBs were maintained on an orbital shaker at 80 rpm from D0 of differentiation.

On D10 of differentiation, EBs were plated onto Matrigel- and laminin- (5 µg/mL; Sigma) coated plates. At D15 of differentiation, cells were considered V-NPCs, as assessed by marker expression after our prior work, were replated using Accutase, and were cultured for a further 2 days in VSM before collection for immunofluorescence. For terminal differentiation, neurospheres were formed from MGE-like progenitors using AggreWell™800 Microwell Culture Plates in ventral telencephalic neuroectoderm differentiation media (VDM) supplemented with 10 µM Y-27632. VDM contained Neurobasal-A (Life Technologies), B-27 supplement (without Vitamin A; Life Technologies), Glutamax (Life Technologies), β-Mercaptoethanol (Life Technologies), Non-Essential Amino Acids (Life Technologies) and Penicillin-Streptomycin Solution (Life Technologies). Neurospheres were maintained for 2 days in AggreWell™800 Microwell Culture Plates and then on an orbital shaker for 2 days at 80 rpm. After 19 days of differentiation from hESCs, neurospheres were plated onto Matrigel- and laminin- (5 µg/mL) coated plates and maintained in VDM supplemented with BDNF (20 ng/mL; PeproTech), with additional medium added 24 hours after plating.

From day 20-35 of differentiation, cultures were fed every 2 days by replacing half of the spent media. On days 22 and 24, media was supplemented with BDNF (20 ng/mL) and CultureOne™ Supplement (Life Technologies), after which media was supplemented with 200 µM ascorbic acid (Sigma-Aldrich), 20 ng/mL BDNF, 10 µM DAPT (Selleckchem), and 2 µM PD0332991

(Selleckchem) until day 30 of differentiation at which point cells were considered V-INs. From day 30-34, VDM was supplemented with 20 ng/mL BDNF, 200 µM ascorbic acid, 200 µM cAMP (Sigma-Aldrich), and 0.5 µM Cytarabine (Tocris Biosciences). For further maturation V-INs were maintained in VDM supplemented with 20 ng/mL BDNF, 200 µM ascorbic acid and 200 µM cAMP, with media changes performed every 3 days by replacing ½ spent media with fresh media.

**Dorsal telencephalic progenitor specification and cortical glutamatergic neuron differentiation**

For specification of hESCs as dorsally patterned neuronal progenitors, EBs were formed as previously described and were then transplanted into dorsal telencephalic neuroectoderm specification media (DSM) supplemented with 10 µM SB-431542, 100 nM LDN-193189, 10 µM Y-27632 and 2 µM XAV-939, constituting D0 of differentiation. DSM contained two-thirds DMEM/F12 (Gibco), one third Neurobasal™ Medium (Gibco), N2 supplement (Life Technologies), B-27 supplement (without Vitamin A), Glutamax, β-Mercaptoethanol, Non-Essential Amino Acids and Penicillin-Streptomycin Solution. EBs were fed on D2 and D4 with DSM supplemented with 10 µM *SB-431542, 100 nM LDN-193189 and 2 µM XAV-939, and from D6 to D10* with DSM supplemented with 10 µM SB-431542 and 100 nM LDN-193189. EBs were plated onto Matrigel- and laminin- (5 µg/mL) coated plates on D10 and were fed with un-supplemented DSM until D20. Cells were collected on D20 as dorsally patterned neuronal progenitors (D-NPCs).

For terminal differentiation, cells were plated onto laminin and poly-D-lysine coated cultureware and fed with dorsal telencephalic neuroectoderm differentiation media (DDM) containing two-thirds DMEM/F12 (Gibco), one third Neurobasal™ Medium (Gibco), N2 supplement (Life Technologies), B-27 supplement, Glutamax, β-Mercaptoethanol and Penicillin-Streptomycin Solution. From day 24-27, DDM was supplemented with CultureOne™ supplement. From D28-32, DDM was supplemented with 5 µM DAPT and 1 µM PD0332991 (PD). Cells were maintained in DDM supplemented with 20 ng/mL BDNF, 200 µM ascorbic acid, and 0.5 µM Cytarabine until D40, at which point they were considered immature glutamatergic neurons (D-INs).

**Direct hPSC differentiation to cortical glutamatergic-like neurons (iGluts)**

Induced neurons were generated as previously described (Schafer et al. 2019). Briefly, hPSCs carrying a construct for inducible expression of Neurogenin-2 (NGN2) were maintained under selection with 0.5 µg puromycin. To begin differentiation, hPSC models were dissociated with Accutase and seeded onto Matrigel-coated cultureware in StemFlex media supplemented with Y-27632 and puromycin and allowed to recover for 48 hours. Expression of NGN2 was then induced by addition of 4 µg/mL Doxycycline (Sigma-Aldrich), constituting day 0 (D0) of differentiation. Cultures were maintained with daily feeding from D0-D6 in StemFlex media (D0-D2) and then in iGlut Media (D3-6) supplemented with Y-27632 and puromycin. iGlut Media was comprised of two-thirds DMEM/F12, one third Neurobasal™ Medium, N2 supplement, B-27 supplement, Glutamax, β-Mercaptoethanol, non-essential amino acids and Penicillin-Streptomycin Solution. From D6-8, cultures were fed daily with iGlut Media supplemented with cytarabine (1µM) and subsequently cultures were fed every 2 days with iGlut Media supplemented with 10 ng/mL BDNF. On D14, cells were harvested as iGluts.

**Dorsal or ventral telencephalic patterning of cortical organoids**

Dorsally or ventrally patterned neurospheres as derived during the specification of D- or V-NPCs were maintained on an orbital shaker for 24 hours instead of being plated on D10. Neurospheres were then embedded in Matrigel© (Corning) using organoid embedding sheets (Stem Cell Technologies) for 30 minutes before being transferred to ultra-low attachment multi-well plates (Corning) in DSM (D-Orgs) or VSM (V-Orgs). 48 hours after embedding media was changed to organoid differentiation media (ODM) containing Neurobasal media supplemented with B-27 supplement (without Vitamin A), Glutamax, N2 supplement, β-Mercaptoethanol, Non-Essential Amino Acids and Penicillin-Streptomycin Solution, insulin solution (3 µg/mL, Sigma-Aldrich), EGF (20 ng/mL, PeproTech) and bFGF (20 ng/mL, PeproTech). Organoids were fed every 48 hours by replacement of half spent media with fresh ODM until D21 when media was changed to organoid maturation media (OMM) containing Neurobasal media supplemented with B-27 supplement (without Vitamin A), Glutamax, N2 supplement, β-Mercaptoethanol, Non-Essential Amino Acids and Penicillin-Streptomycin Solution, insulin solution (3 µg/mL, Sigma-Aldrich), cAMP and AA. Organoids were then fed every 72 hours by replacement of half spent media with fresh OMM until collection on D30.

**Immunocytochemistry**

Organoids were fixed overnight at 4°C in 4% Paraformaldehyde (Sigma-Aldrich) before being cryopreserved in 30% (v/v) sucrose solution (Sigma-Aldrich) overnight at 4°C. Organoids were them embedded in a 1:1 solution of O.C.T. and 30% sucrose before being cryosectioned at 8 µm on a leica CM1510 cryostat. Antigen retravel was performed on organoid sections using a citrate buffer (Sigma-Aldrich) at 80°C for 12 minutes. Non-specific antibody binding was blocked by incubating sections with 5% donkey serum in tris-buffered saline (TBS) with 1% Triton X-100 for 1 hour at room temperature. Primary antibodies (Supplementary Table 2) were added overnight at 4°C in TBS with 0.5% Triton X-100. Sections were washed thrice with TBS being secondary antibodies (Supplementary Table 2) were applied for 1 hour at room temperature in TBS. Sections were again washed thrice in TBS before being counterstained with DAPI and mounted using ProLong™ Glass Antifade Mountant.

D- or V-NPCs were passaged using Versene (Gibco) onto Matrigel and laminin coated plasticware at a ratio of 1:1.5 and were maintained in the corresponding media DSM for D-NPCs or VSM supplemented with SB-431542, LDN-193189, SAG and XAV-939 for V-NPCs. Medias were supplemented with Y-27632 for the passage and then cells were fed with corresponding medias 48 hours after passaging by half media replacement. Cells were fixed 4 days after passaging in 4% Paraformaldehyde before blocking in phosphate buffer saline (PBS) containing 5% donkey serum and 0.1% Tween20. Primary antibodies (Supplementary Table 2) were added overnight at 4°C in PBS with 0.1% Tween20. Cells were washed with PBS and incubated with secondary antibodies (Supplementary Table 2) in PBS for 1 hour at room temperature. Cells were counterstained with DAPI and mounted using ProLong™ Glass Antifade Mountant.

For staining of V-INs, cells were passaged as single cells using Accutase on day 28 of differentiation at a density of $2x10^5$ cells/$cm^2$ onto poly-ornithine and laminin coated cover slips. Cells were maintained as previously described until either day 40 or 50 of differentiation and were then fixed using 4% Paraformaldehyde before blocking in phosphate buffer saline (PBS) containing 5% donkey serum and 0.1% Tween20. Primary antibodies (Supplementary Table 2) were added overnight at 4°C in PBS with 0.05% Tween20. Cells were washed with PBS and incubated with secondary antibodies (Supplementary Table 2) in PBS for 1 hour at room temperature. Where appropriate cells were counterstained with DAPI and all cells were mounted using ProLong™ Glass Antifade Mountant.

Images were obtained using a custom-built spinning disc confocal from Intelligent Imaging Innovations (3i) equipped with a phasor for precise photomanipulation, a Zeiss LSM 980 with Airyscan 2 or a Zeiss AxioScan 7 and images were processed with ImageJ or Zeiss Zen software. Quantification was performed using a minimum of 3 random images taken from 3 independent biological replicate experiments for all adherent cultures or using images of whole organoids utilizing 4-9 organoids per batch with a minimum of independent batches generated per biological group. Quantifications were performed using CellProfiler (Stirling et al. 2021), requiring a minimum of 60% overlap between immunofluorescence signal and DAPI staining for co-localization to be scored or using Intellicount (Fantuzzo et al. 2017) to quantify synapse density.

**Reverse Transcription and Quantitative PCR (RT-qPCR)**

RNA was isolated using the NucleoSpin RNA kit (Macherey-Nagel) or TRIzol Reagent (Invitrogen), with ≥1 µg used to make cDNA using the High-Capacity cDNA Reverse Transcription Kit. Equal quantities of cDNA were used as template for quantitative PCR (qPCR), using

PowerTrack™ SYBR Green Master Mix (Applied Biosystems) and the Applied Biosystem StepOne Plus quantitative PCR system. GAPDH was used as an endogenous quantitative control and all primers used can be found in Supplementary Table 3. Data was generated from a minimum of 3 biological replicate samples, each of which consisted of 3 technical replicates. All statistics were performed using abundance relative to endogenous control, and data presented as fold change by comparison with appropriate control.

**Western Blotting**

Protein was isolated in RIPA buffer with Halt™ Protease and Phosphatase Inhibitor Cocktail and western blotted using Bolt™ 4 to 12%, Bis-Tris, 1.5 mm, Mini Protein Gels and then transferred to 0.2 µm nitrocellulose membrane. Nonspecific antibody binding was blocked by 5% milk solution, and primary antibodies (Supplementary Table 2) were incubated with free floating membranes overnight at 4°C. Secondary antibodies were HRP conjugated Goat anti-Rabbit IgG and Goat anti-Mouse IgG, each used at a 1:10,000 dilution. Visualization was performed by chemiluminescence using SuperSignal™ West Pico PLUS Chemiluminescent Substrate, all band intensities were normalized to GAPDH loading controls, and a minimum of 3 biological replicates were averaged before statistical analysis. Statistical analysis was performed on normalized signal intensity, with data presented as fold change compared to appropriate controls. All unmodified western blot images are presented in Supplementary Figure S16.

**Neurosphere Size and Outgrowth Measurements**

Images were taken from D12 of differentiation, using a 4x objective, for 20 neurospheres per biological replicate across a minimum of 3 biological replicates per condition. Sphere size and outgrowth were measured using ImageJ and outgrowth is defined as the area of cells outgrown from the plated neurosphere, normalized to corresponding sphere size.

**Manipulation of EZH2 expression and function**

D-NPCs were replated using Versene onto Matrigel and laminin coated plasticware at a 1:1.5 ratio in DSM supplemented with Y-27632. For pharmacological EZH2 inhibition the EZH2 inhibitor GSK343 (R&D Systems) was applied 24 hours after passaging at a concentration of 4 µM and compared to cells treated with 0.04% DMSO. Cells were fed by half media change after a further 2 days and 4 days after passaging with DSM supplemented with either GSK343 or DMSO and cells were collected 5 days after passaging for further analysis. For siRNA mediated knockdown of EZH2, D-NPCs were again passaged using Versene onto Matrigel and laminin coated plasticware at a 1:1.5 ratio in DSM supplemented with Y-27632 and then 5 pmol of siRNAs (Supplementary Table 4) targeting EZH2 or GFP were applied using Lipofectamine® RNAiMAX Regent (Invitrogen) as per the manufacturer's instructions after 24 hours of recovery. Cells were then maintained for a further 4 days in DSM with half media changes on day 1 and 3. Cells were then collected for down stream analysis.

For EZH2 overexpression experiments V-NPCs were passaged using Accutase onto Matragel and laminin coated plasticware at a density of $1x10^5$ cells/$cm^2$ in VSM supplemented with SB-431542, LDN-193189, SAG and XAV-939. After 24 hours of recovery, cells were transduced with pHAGE-EZH2 lentivirus with the aid of polybrene. Cells were then maintained for the subsequent 4 days with media replaced on days 1 and 3 using VSM supplemented with SB-431542, LDN-193189, SAG and XAV-939 before cells were collected for further analysis.

**RNA-seq analysis**

RNA was isolated using the AllPrep DNA/RNA kit (Qiagen) and RNA-seq library preparation was performed using the SMARTer Ultra Low RNA kit (Takara-Clontech). Reads were obtained using an Illumina NovaSeqX Plus sequencer, to a depth of ~30 million (M) reads per sample by the Washington

University Genome Technology Access Center at McDonnell Genome Institute (GTAC@MGI). Raw reads from RNA-seq samples were quality trimmed using cutadapt (v2.4)(Kechin et al. 2017) with the options of quality-cutoff=15,10 and minimum-length=36. Reads were aligned and quantified using hg38 GENCODE V42 comprehensive gene annotations using salmon(Patro et al. 2017) in mapping based mode and correcting for sequence and GC bias (--seqBias --gcBias) and options (--validateMappings and --rangeFactorizationBins 4). Quantifications were summarized to the gene level using tximport (v1.32)(Soneson et al. 2015) using hg38 GENCODE V42 comprehensive gene annotations.

For differential expression analysis, only genes with >5 counts in >2 samples were included. Differential expression analysis was performed using DESeq2(Love et al. 2014) in negative binomial mode, using counts for genes that passed the cutoff. A 1.5-fold linear expression change and a Benjamini and Hochberg false discovery rate of < 0.05 (FDR; adjusted p-value) were set as cutoff values for significance. Raw results from each DESeq2 analysis are presented as supplementary data. Expression data is presented and visualized using transcript per million mapped reads (TPM). Raw results are available from GEO under accession GSE294189.

**Whole Genome Bisulfite Sequencing (WGBS)**

DNA was isolated using the AllPrep DNA/RNA kit and 40-300ng of DNA was fragmented for 45 seconds using the Covaris E220 sonicator (10% Duty Factory, 175 Peak Incidence Power, 200 cycles per burst, milliTUBE 200µL AFA Fiber) and purified using 0.7 volumes of SPRISelect Beads. A small amount of unmethylated lambda DNA was added for non-conversion rate quality control, and DNA was bisulfite converted using the EZ DNA Methylation-Direct Kit. Libraries were generated using the xGen™ Methyl-Seq Lib Prep Kit with combinatorial dual indexes as instructed, using 10-15 cycles of final amplification. Libraries were pooled and sequenced at the Washington University GTAC@MGI on the NovaSeq 6000 2x150.

Paired end sequencing reads were trimmed to remove adapters and low-quality sequence using TrimGalore (v0.6.7, --paired --clip_R1 10 --clip_R2 10). Trimmed reads were then mapped to the human genome (hg38) using bismark (v0.23.1, default parameters: -q --score-min L,0,-0.2 -p 4 --reorder --ignore-quals --no-mixed --no-discordant --dovetail --maxins 500 --directional). Aligned reads were filtered based on mapping quality (>=10) using Samtools (v0.1.19) and PCR duplicates were removed using GATK/Picard MarkDuplicates (gatk v4.1.3.0). Methylation data at individual cytosines was extracted using MethylDackel extract (v0.6.01, with --CHG and –CHH flags and otherwise default parameters). Methylation read counts were merged across strands using a custom python script. DMRs were computed using DSS (v2.48.0) using a minimum of 3 independent biological replicates per condition. The test function DMLtest was run with smoothing=TRUE and otherwise default parameters. DMRs were then computed with a *p*-value threshold of 0.01. DMRs have a minimum length of 50 bp, must contain 3 or more CpGs, and DMRs within 50 bp are merged (DSS default parameters). DMRs were annotated using ChIPseeker (v1.36.0, with the tssRegion set to +/- 3000 bp and otherwise default settings) and the TxDb.Hsapiens.UCSC.hg38.knownGene human gene annotations. Genome browser tracks were visualized using bedGraph files and the WashU Epigenome browser. All raw and processed data is available from dbGaP (in process) or GEO (GSE294191).

**CUT&Tag**

CUT&Tag was performed on all samples as previously described (Chapman et al. 2024) specifically using 2-5x$10^5$ cells per sample. Primary and secondary antibodies used for CUT&Tag in this study can be found in Supplementary Table 2, with a minimum of 4 independent biological replicates used for each time point and sample set. Samples with unique dual-end indexes were pooled in equimolar concentrations and sequenced by GTAC@MGI on the NovaSeq-6000 sequencer as 150 bp paired-end reads at a depth of ≥5 million (M) reads per sample. Raw reads from CUT&Tag samples were processed by AIAP (v1.1) (Liu et al. 2021), using human genome hg38 as a reference to perform read quality control, alignment, quantification, and peak calling.

Analysis of differential binding from CUT&Tag data was performed using DiffBind (Ross-Innes et al. 2012), using trimmed and aligned binary alignment and map files along with peaks identified by MACS2 within the AIAP (v1.1) workflow. Blacklisted sequences were removed and a minimum overlap of 3 samples was used to generate the consensus peak set for each biological condition. Read depth data was normalized using the reads in peaks library normalization method and significantly differentially bound peaks were identified using DESeq2, with a Benjamini and Hochberg FDR of <0.05. Complete result tables for significant peak changes in signal for each comparison are presented as Supplementary Data. Peak sets, including differentially bound peaks, were annotated using ChIPseeker (using the same parameters as WGBS analysis). Curated peak sets were visualized using deepTools (Ramirez et al. 2016), using the computeMatrix reference-point and plotHeatmap functions. All raw and processed data is available from GEO (GSE294186).

**Neuronal Morphology**

Immature D28 cortical interneurons were replated using Accutase as previously described onto poly-d-ornithine and laminin coated cultureware at a density of $2x10^4$ cells per $cm^2$ in VDM supplemented with 2 µM Y-27632. After 24 hours recovery media was replaced with fresh VDM supplemented with 20 ng/mL BDNF. After a further 24 hours neurons were imaged using a standard phase-contrast setup at 10x magnification with 4 random images taken from each biological replicate. Neuronal morphology was manually traced using ImageJ, quantifying morphometric parameters from 10 neurons per image for a total of 40 neurons per biological replicate. Example traces are each taken from separate biological replicates and are selected as representative of the morphology observed.

**Electrophysiological Recording Parameters**

Recording electrodes were filled with (in mM): 140 K-glucuronate, 10 NaCl, 5 $MgCl_2$, 0.2 EGTA, 5 Na-ATP, 1 Na-GTP, and 10 HEPES, with the pH adjusted to 7.4 with KOH. Open-tip resistance was 2-6 MOhm. GABAergic or glutamatergic neuronal cultures were perfused with Tyrode's solution (in mM): 150 NaCl, 4 KCl, 2 $MgCl_2$, 2 $CaCl_2$, 10 Glucose, 10 HEPES, pH adjusted to pH 7.4 with NaOH. Whole-cell currents and membrane potentials were recorded with an Axopatch 200A amplifier (Molecular Devices). Cell capacitance and input resistance as well as peak inward sodium current and steady-state outward potassium current during depolarizing voltage steps from a holding potential of -80 mV were determined from voltage clamp recordings(Meganathan et al. 2017). Current clamp recordings, spontaneous inhibitory synaptic currents, and currents evoked by 100 µM gaba-aminobutyric acid (GABA) and kainate were obtained in a modified extracellular solution (in mM): 120 NaCl, 3 KCl, 10 glucose, 1 $NaH_2PO_4$, 4 $NaHCO_3$, 5 HEPES, pH adjusted to 7.4 with NaOH, which was delivered from an 8-barrelled local perfusion pipette positioned near the recorded cell(Meganathan et al. 2017).

**Low Density MEA Preparation and Recording**

D24 V-INs and D28 D-INs were dissociated and combined, resulting in a 70:30 ratio of D:V INs, and plated onto laminin- and poly-D-lysine-coated cultureware. Cells where then maintained in DDM supplemented with 20 ng/mL BDNF, 200 µM ascorbic acid and 0.5 µM Cytarabine for a further 7 days, after which cell cultures were fed with maturation media (MM) comprised of BrainPhys™ Neuronal Medium (Stem Cell Technologies) supplemented with B27 plus supplement (Gibco), penicillin, streptomycin, 10 ng/mL BDNF and 200 µM ascorbic acid. 17 days after re-plating, cells were dissociated and were plated as high density drop cultures (5,000 cells/µL) containing 20 µg/ml laminin onto MEAs pre-treated with 0.01% polyethylenimine (Sigma). After 1-hour, conditioned MM was added into each MEA and after 24 hours 0.5 mL of fresh MM was added to each array.

MEA cultures were maintained in MM replaced every 3-4 days. Electrophysiological activity was recorded every 5 days using hardware (Maestro Edge) and software (AxIS 1.5.2) from Axion Biosystems. Channels were sampled simultaneously with a gain of 1000× and sampling rate of 12.5 kHz/channel. During the recording, constant temperature was maintained at 37°C, and the chamber was maintained at 5% $CO_2$. A Butterworth band-pass filter (with a high-pass cut-off of 200 Hz and low-pass cut-off of 3000Hz) was applied along with a variable threshold spike detector set at 4.5× standard deviation on each channel. Synchronized bursts (SBs) were identified using an envelope algorithm, defining a SB by identifying periods when activity exceeds a threshold of 1 x standard deviation above the mean with a minimum of 200ms between SBs and at least 35% of electrodes included. A minimum of 4 MEAs per cell line per differentiation (technical replicates) have been considered for the analysis with a minimum of 3 separate differentiations used per time point.

**High Density MEA Preparation and Recording**

MEAs were cleaned and prepared by sequentially washing electrode with ethanol and DPBS before sterilization was performed by 15-minute exposure to UV. MEAs were simultaneously coated with poly-ornithine and polyethylenimine overnight at 37°C before they were washed with water and dried at room temperature in a sterile environment. D6 iGlut differentiations and D28 V-IN differentiations were dissociated and combined in a 70:30 ratio of iGlut:V-INs and this suspension was then adjusted to produce a 1,500 neuron/µL suspension containing 40 µg/ml laminin and 250 astrocytes/µL using rat primary cortical astrocytes (Gibco™). The resulting solution was plated directly onto each MEA (using 25 µL per MEA) and then MEAs were incubated for 1.5 hours at 37°C before fresh MM was added to each MEA. MEA cultures were maintained in MM replaced every 3-4 days. Electrophysiological activity was recorded from 2304 channels sampled simultaneously from each MEA at a rate of 10 kHz/channel with raw data high-pass filtered at 100 Hz and acquired with noise blanking compression (3.5 standard deviations (SD)/4 SD low/high threshold). The recording chamber was maintained between 36.5 and 38°C and 5% $CO_2$ for the duration of each 10-minute recording. Action potentials (spikes) were defined using a precise timing spike detection algorithm as activity with greater amplitude than 7x standard deviation on each channel. An electrode burst was defined using an inter-spike-interval (ISI) algorithm, as a period with a minimum of 10 spikes separated by a maximum ISI of 70 ms and a bursting electrode was defined as an electrode with at least 2 bursts in a 10-minute recording. Finally neuronal network bursting was defined using a recruitment-based algorithm with a minimum threshold of 0.1 spikes/s across a minimum of 30% of active electrodes, using a 150 ms bin size, and requiring a 100-spike minimum. All active electrodes (spiking rate > 0.1 Hz) were included in the analysis of neuronal and neuronal network activity unless otherwise stated. For each condition 3 MEAs were plated with the same MEAs recorded longitudinally.

### Supplementary Tables

#### Supplementary Table 1: Guide RNA sequences

| NAME | SEQUENCE |
|---|---|
| **DNMT3A G1** | GGCGGCGGCGGCGAGAGCAG |
| **DNMT3A G2** | GACGCGGCGCCGCGGCACCA |

#### Supplementary Table 2: Antibodies

| TARGET | SPECIES | SUPPLIER | CATALOG NUMBER |
|---|---|---|---|
| **KI67** | Rabbit | Abcam | ab15580 |
| **H3K27ME3** | Rabbit | Cell Signaling | 9733 |
| **DNMT3A** | Rabbit | Abcam | ab2850 |
| **GAPDH** | Mouse | Cell Signaling | 97166 |
| **DLX2** | Rabbit | Invitgrogen | MA5-47229 |
| **NKX2.1** | Rabbit | Cell Signaling | 12373S |
| **SOX2** | Rabbit | Cell Signaling | 30064S |
| **SST** | Rabbit | Invitrogen | 701935 |
| **TBR1** | Rabbit | Abcam | ab183032 |
| **TBR2** | Rabbit | Abcam | ab216870 |
| **TUJ1** | Mouse | Cell Signaling | 4466S |
| **ASCL1** | Rabbit | Abcam | ab211327 |
| **MTOR** | Rabbit | Cell Signaling | 2983S |
| **P-MTOR** | Rabbit | Cell Signaling | 5536S |
| **AKT** | Rabbit | Cell Signaling | 4691S |
| **P-AKT** | Rabbit | Cell Signaling | 9275S |
| **RPS6** | Rabbit | Cell Signaling | 2217S |
| **P-RPS6** | Rabbit | Cell Signaling | 5364S |
| **CALB1** | Rabbit | Cell Signaling | 13176S |
| **SYN1** | Rabbit | Synaptic Systems | 106008 |
| **PSD95** | Mouse | Synaptic Systems | 124011 |
| **TUJ1** | Guinea Pig | Synaptic Systems | 302304 |
| **ANTI RABBIT IGG** | Goat | Cell Signaling | 35401S |
| **ANTI MOUSE ALEXA FLOUR 488** | Donkey | Invitrogen | A21202 |
| **ANTI RABBIT ALEXA FLUOR PLUS 555** | Donkey | Invitrogen | A32794 |
| **ANTI GUINEA PIG ALEX FLOUR 555** | Goat | Invitrogen | A21435 |
| **ANTI MOUSE ALEXA FLOUR PLUS 405** | Donkey | Invitrogen | A48257 |
| **ANTI RABBIT ALEXA FLUOR PLUS 488** | Donkey | Invitrogen | A32790 |

#### Supplementary Table 3: qPCR Primers

| TARGET | FORWARD PRIMER | REVERSE PRIMER |
|---|---|---|
| **FGFR2** | GGAAAGTGTGGTCCCATCTGA | TCCAGGTGGTACGTGTGATTG |
| **LHX5** | TGTACGGACCGCAGTTTGTC | CCGACGAGGTCGAGTTGTC |
| **PAX6** | TGGGCAGGTATTACGAGACTG | ACTCCCGCTTATACTGGGCTA |
| **POU3F2** | AAGCGGAAAAAGCGGACCT | GTGTGGTGGAGTGTCCCTAC |
| **WNT7B** | CACAGAAACTTTCGCAAGTGG | GTACTGGCACTCGTTGATGC |
| **MEI4** | TCCCGTGCCCGCTTGAG | CATCCTGGCTTTTGTCCCTGAAAA |

| | | |
|---|---|---|
| **BARHL1** | GGGATCGACTCCATTCTCTCC | GTCCCCGTCTCCAAACAGT |
| **DMRT3** | ATGTGGCAAAGAGTAAGGGCT | GCGGTCTGTTGGCTTTCAAG |
| **NHLH2** | GCAGCAGATTCGGACCATCC | ACGTGGTTGAGATAGGAGATGT |
| **ZNF536** | CCTGGGATGCCTCAATCTCG | AAGCCTGGTAGCTGTTCACG |
| **CDH23** | CGCCCACATTTCACAATCAGC | CGTCCCCACTGGTGTATTCT |
| **DRD4** | ATCAGCGTGGACAGGTTCGT | CACCCTGCCGGTTGTAGC |
| **DSCAML1** | TTTTCAGGGAACCCTACACCG | ACGCTAACATATTCCTGCACTG |
| **FAM107A** | GCAGCGTGTCCTAGAGCAC | CCGCAGGTTTTCCCTGACT |
| **GRM1** | CCTCTGTATCGCCCATTCTGA | GGAAGCCTCTCTCGGAGTTT |
| **IL31RA** | AACATAGCGAAAACTGAACCACC | GCCAACTCAGGCTTTATCCATTC |
| **KCNIP1** | CAGGTCCTTTATCGAGGCTTCA | CGACAGAGCGGTTACAAAGTC |
| **MCHR1** | GTGGCTGTATGCCAGACTCAT | CAGGGTGAACCAGTAGAGGTC |
| **MYO7A** | GCAGCACTCGTGGATTGAG | TGTACTTTCCGAAACGGCTTG |
| **NEU4** | GGCCACGGGATGACAGTTG | CAGGCGGATACCCATGTGTAG |
| **ADGRL3** | GCCTATGTCCAGGCAATGGT | AAGCTTGTGGCTGAAGGAGG |
| **CABP1** | GAGAGGCTATGAGGAAGCTCC | TGAGGTCCACATCTCGGATAAT |
| **EPHA3** | ACTCTACGAGACTGCAATAGCA | TCCCCAAGATCCATTTGAGTGA |
| **GRIA1** | CGAGCTTTCCCGTTGATACAT | TCTGCCACTTGTAATGGTCAATG |
| **GRIN2A** | TCATGCAGGATTATGACTGGCA | TGTGGTCTTGACGAAGCTGAT |
| **GRIP2** | GCAGGGGAGACAATAGCGAAC | CACAGTGATCCCTCGGAACT |
| **LRRC4B** | TCGACGACCTCAAGTCGCT | GGGCGCATAGCAGGTGAAAT |
| **NTRK2** | ACCCGAAACAAACTGACGAGT | AGCATGTAAATGGATTGCCCA |
| **PCDHGC3** | TGGCCCAACAACCAGTTTGA | CCATCAGCAGCTTCACTGGC |
| **SEMA3A** | CTATCTTCCGAACTCTTGGGCA | CTTTGGATCATTGAGCCACCT |
| **SLITRK2** | CCAACGCGGTGACTCTTCA | CCTCGATGGCACTGATGTAATTG |
| **TPBG** | TGAGCCTGACCTACGTGTC | GCCATTGTGAAGGACCTTGAG |

**Supplementary Table 4: siRNA sequences**

| **NAME** | **IDT ITEM NAME** | **FORWARD SEQUENCE** | **REVERSE SEQUENCE** |
|---|---|---|---|
| **SCRAM** | Negative Control DsiRNA | N/A | N/A |
| **EZH2 SIRNA 1** | hs.Ri.EZH2.13.1 | rGrArGrGrArUrCrArCrCrGrArGrArUrGrArUrArArArGrAAA | rUrUrUrCrUrUrUrArUrCrArUrCrUrCrGrGrUrGrArUrCrCrUrCrCrA |
| **EZH2 SIRNA 2** | hs.Ri.EZH2.13.2 | rCrArUrUrGrGrCrArCrUrUrArCrUrArUrGrArCrArArUrUTC | rGrArArArUrUrGrUrCrArUrArGrUrArArGrUrGrCrCrArArUrGrArG |
| **EZH2 SIRNA 3** | hs.Ri.EZH2.13.3 | rUrArArUrGrCrArGrUrArUrGrGrUrArCrArUrUrUrUrUrCAA | rUrUrGrArArArArArUrGrUrArCrCrArUrArCrUrGrCrArUrUrArUrU |

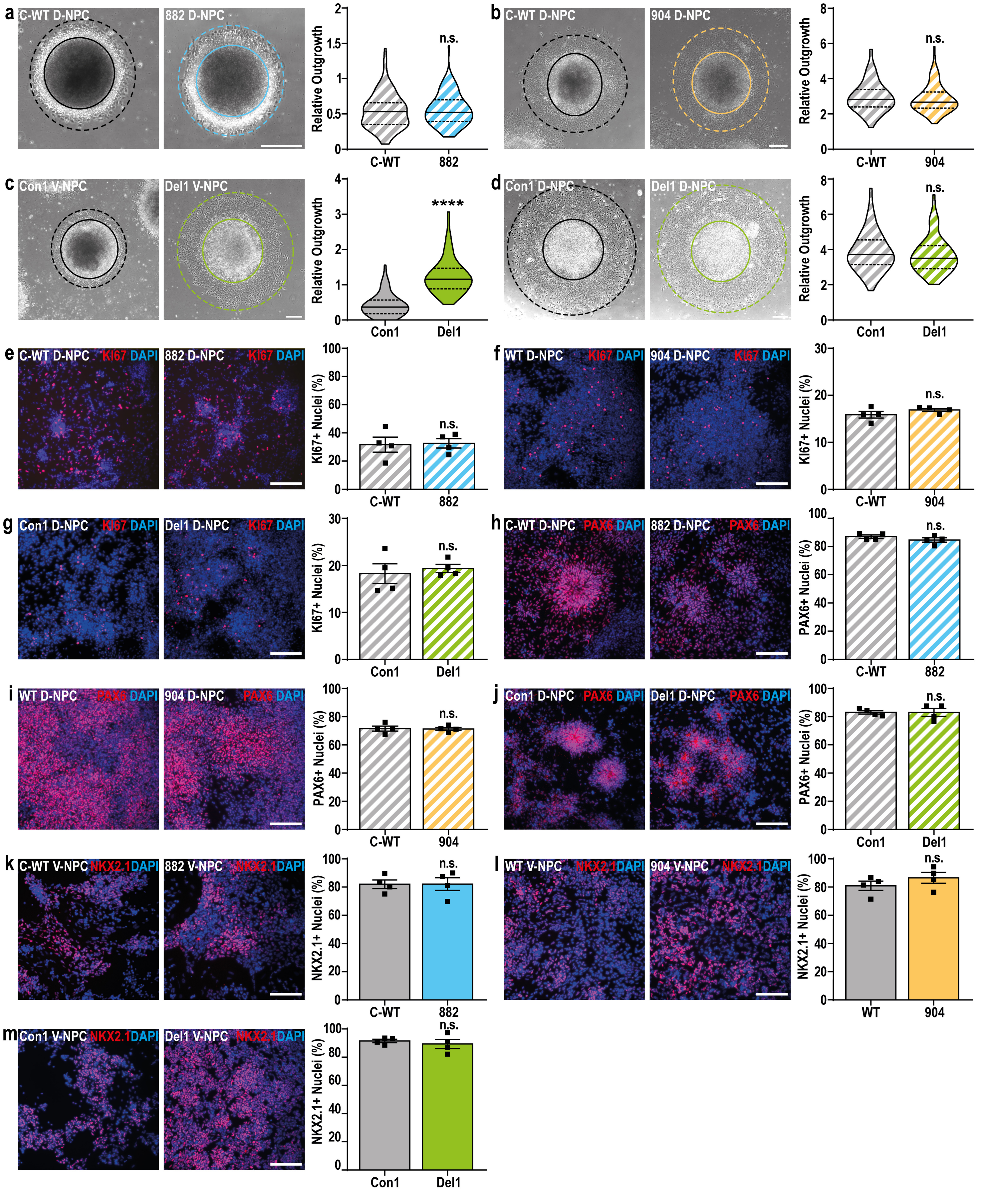


**Supplementary Fig. S1: Characterization of NPC growth and specification in 882, 904 and Del1 NPCs**

**a-d**, Representative images and quantification of neurosphere outgrowth, defined as outgrowth (dotted line) minus neurosphere size (bold line) relative to neurosphere size, in D12 control D/V-NPC versus (**a**) 882 D-NPC, (**b**) 904 D-NPC, (**c**) Del1 V-NPC and (**d**) Del1 D-NPC. **e-g**, Representative images and quantification of the proportion of KI67 positive nuclei in control D-NPCs vs (**e**) 882 D-NPCs, (**f**) 904 D-NPCs and (**g**) Del1 D-NPCs. **h-j** Representative images and quantification of the proportion of PAX6 positive nuclei in control D-NPCs versus (**h**) 882 D-NPCs, (**i**) 904 D-NPCs and (**j**) Del1 D-NPCs. **k-l,** Representative images and quantification of the proportion of NKX2.1 positive nuclei in control V-NPCs versus (**k**) 882 V-NPCs, (**l**) 904 D-NPCs and (**m**) Del1 V-NPCs. Data is represented as (**a-d**) data distributions with median (bold line) and upper and lower quartiles (dotted lines) indicated or as (**e-m**) mean +/- SEM with individual biological replicates indicated. All data was analyzed by Student's t-test, versus matched isogenic controls with n=4 biological replicate experiments for all conditions, n.s.=not significant, pValue>0.05, *pValue<0.05, **pValue<0.01, ***pValue<0.001, ****pValue<0.00001. Scale bar = 100μm.

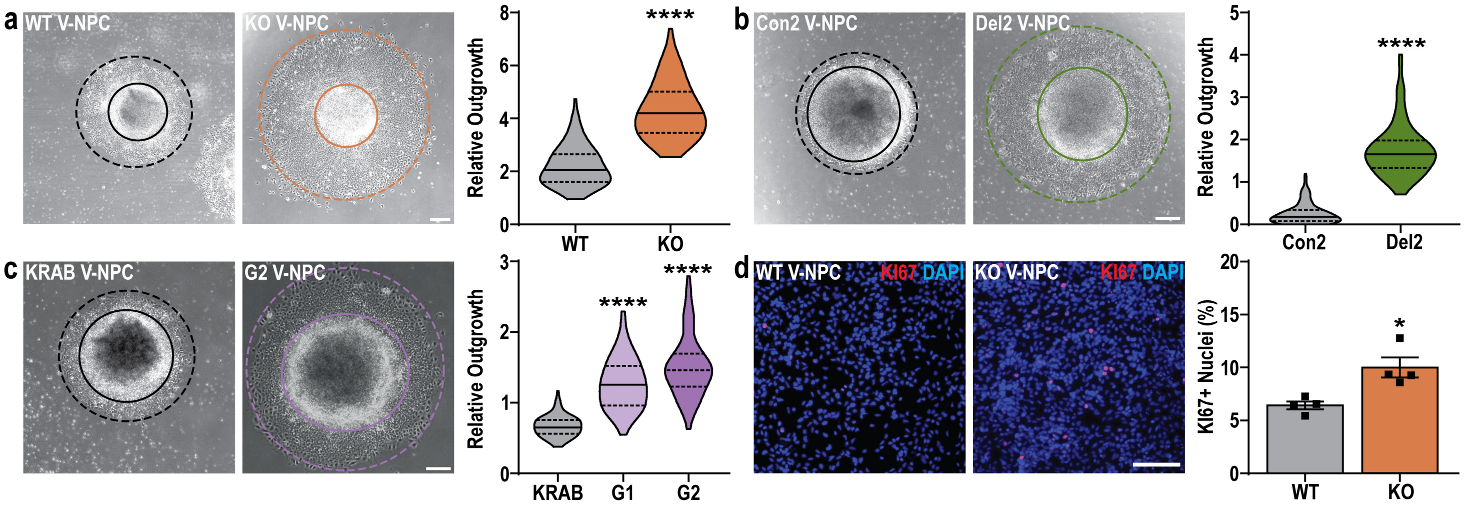


**Supplementary Fig. S2: Characterization of V-NPC growth in Del2, KO and CRISPRi NPCs**

**a-c**, Representative images and quantification of neurosphere outgrowth, in D12 control V-NPCs versus (**a**) KO V-NPCs, (**b**) Del2 V-NPCs and (**c**) G1/G2 V-NPCs. **d**, Representative images and quantification of the proportion of KI67 positive nuclei in control V-NPCs versus KO V-NPCs. Data is represented as (**a-c**) data distributions with median (bold line) and upper and lower quartiles (dotted lines) indicated, or as (**d**) mean +/- SEM with individual biological replicates indicated. All data was analyzed by Student's t-test, versus matched isogenic controls with n=4 biological replicate experiments for all conditions, *pValue<0.05; ****pValue<0.00001. Scale bar = 100 μm.

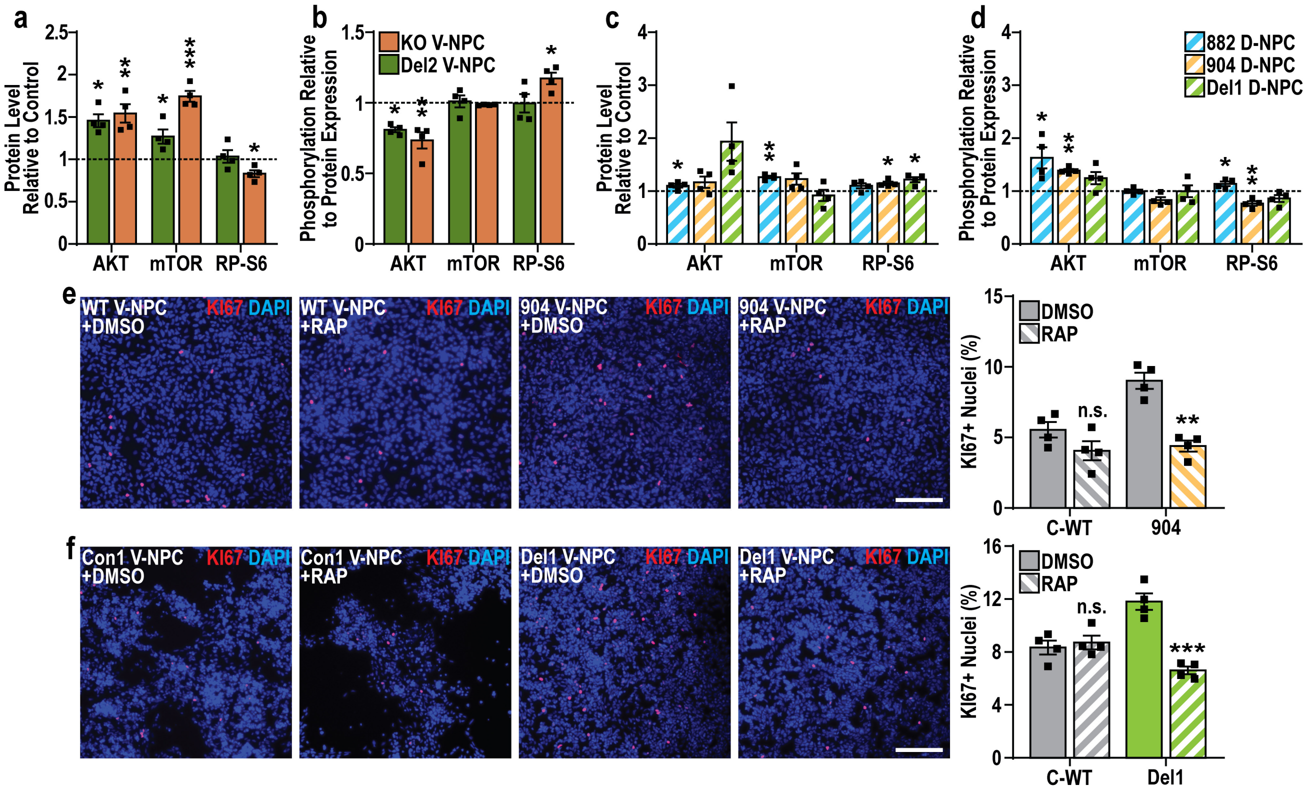


**Supplementary Fig. S3. Effects of DNMT3A LOF on PIK3/AKT/mTOR signaling**

**a-d**, Quantification of relative (**a**,**c**) abundance and (**b**,**d**) phosphorylation of AKT, mTOR and ribosomal protein S6 in (**a**,**b**) TBRS V-NPCs (KO and Del2) or (**c**,**d**) TBRS D-NPCs (882, 904 and Del1) compared to matched controls (indicated by dotted lines). **e-f**, Representative images and quantification of KI67 positivity in (**e**) 904 and (**f**) Del1 V-NPCs compared to matched controls after treatment with either DMSO or rapamycin (RAP). Data is represented as mean +/- SEM with individual biological replicates indicated (**a-f**) and was analyzed by Student's t-test, versus matched controls (**a-d**) or by one-way ANOVA with results shown as comparisons between RAP treated and DMSO treated cells (**e-f**). n=4 biological replicate experiments for all conditions, pValues: *p<0.05;**p<0.01;***p<0.001. Scale bars=100 µm.

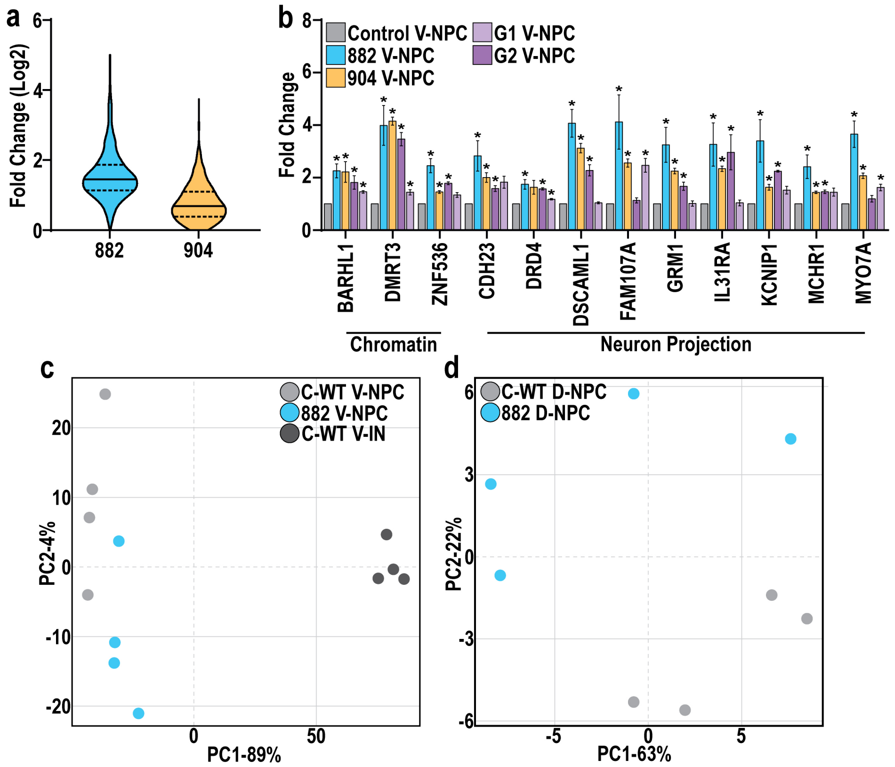


**Supplementary Fig. S4: Extended characterization of transcriptomic changes in 882 and 904 NPCs**

**a**, Magnitude of gene expression changes of shared-DEGs in 882 and 904 V-NPCs versus matched isogenic control V-NPCs. **b**, mRNA expression changes of shared-DEGs associated with 'Chromatin' or 'Neuron Projection' GO terms across 882, 904 and CRISPRi (G1/G2) V-NPCs, compared to matched isogenic control V-NPCs. **c-d,** Principal component analysis comparing (**c**) control V-NPCs and V-INs with 882 V-NPCs and (**d**) control D-NPCs with 882 D-NPCs. Data is represented as (**b**) mean +/- SEM and was analyzed by Student's t-test, versus matched isogenic controls with n=4 biological replicate experiments for all conditions; *pValue<0.05.

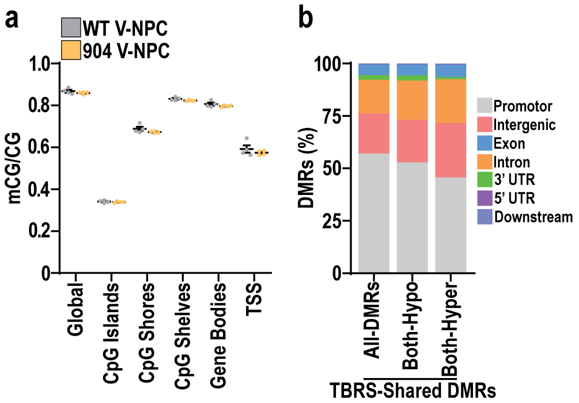


**Supplementary Fig. S5: Extended characterization of mCG changes in 882 and 904 NPCs**

**a**, mCG/CG levels in control (WT) versus 904 V-NPCs. **b,** Genomic location of TBRS shared DMRs, categorized by gene feature and expressed as a percentage of each DMR category. Data is represented as (**a**) mean +/- SEM and was analyzed by Student's t-test, versus matched isogenic control with n=4 biological replicate experiments for all conditions.

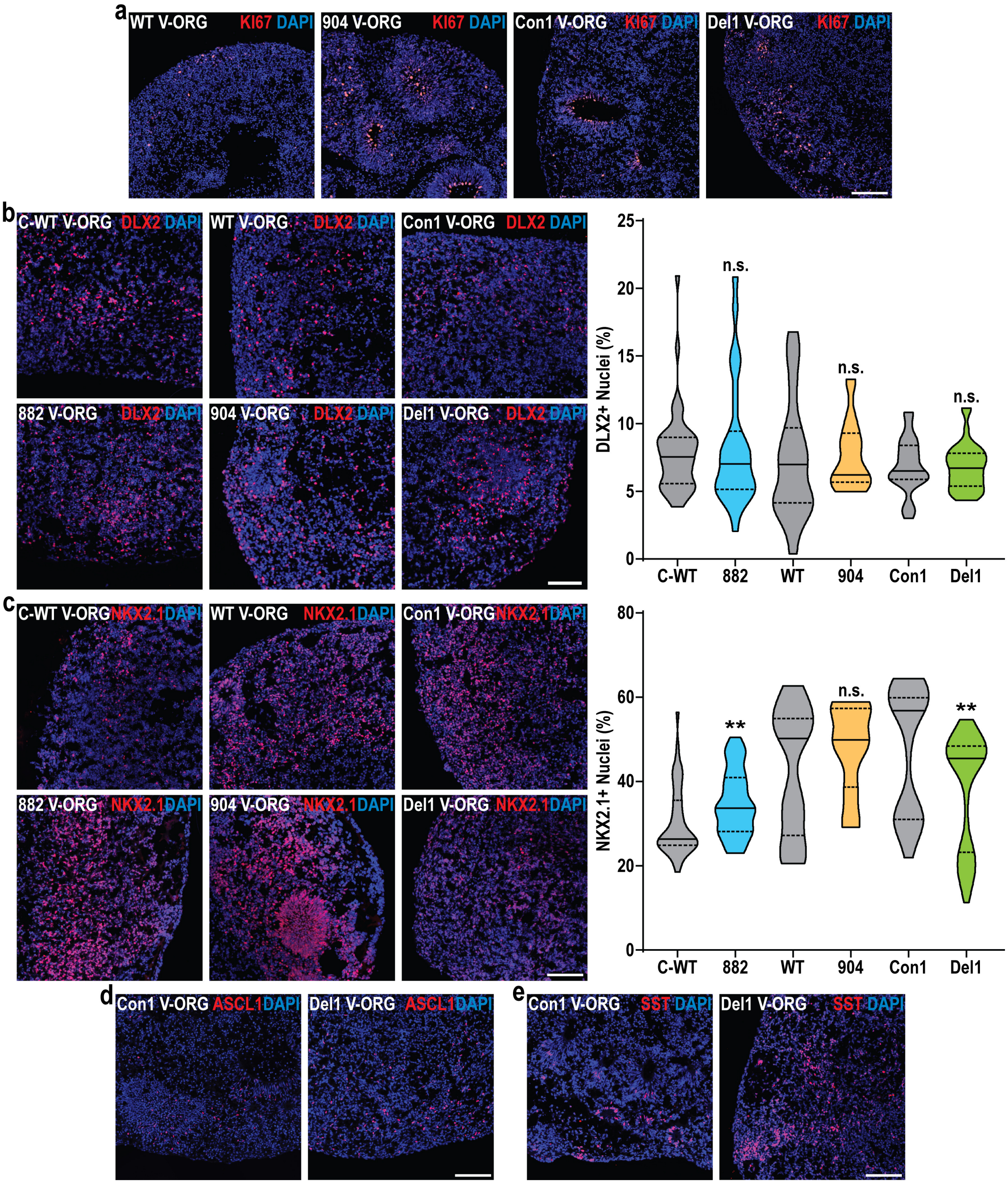


**Supplementary Fig. S6: Changes in proliferation and neurogenesis in TBRS V-ORGs**

**a,** Example images of the proportion of KI67+ nuclei in control (WT and Con1) and TBRS (904 and Del1) V-ORGs. **b-c,** Example images and quantification of the proportion of (**b**) DLX2+ or (**c**) NKX2.1+ nuclei in control and TBRS V-ORGs. **d-e,** Example images of the proportion of (**d**) ASCL1+ or (**e**) SST+ nuclei in Con1 and Del1 V-ORGs. All data is represented as distributions with median (bold line) and upper and lower quartiles (dotted lines) indicated and was analyzed by Mann-Whitney tests comparing each TBRS model to its appropriate isogenic control. A minimum of 3 batches of organoids were prepared for each condition with 3-9 organoids assessed per batch. pValues: **p<0.01; n.s.-non-significant. Scale bars=100 µm.

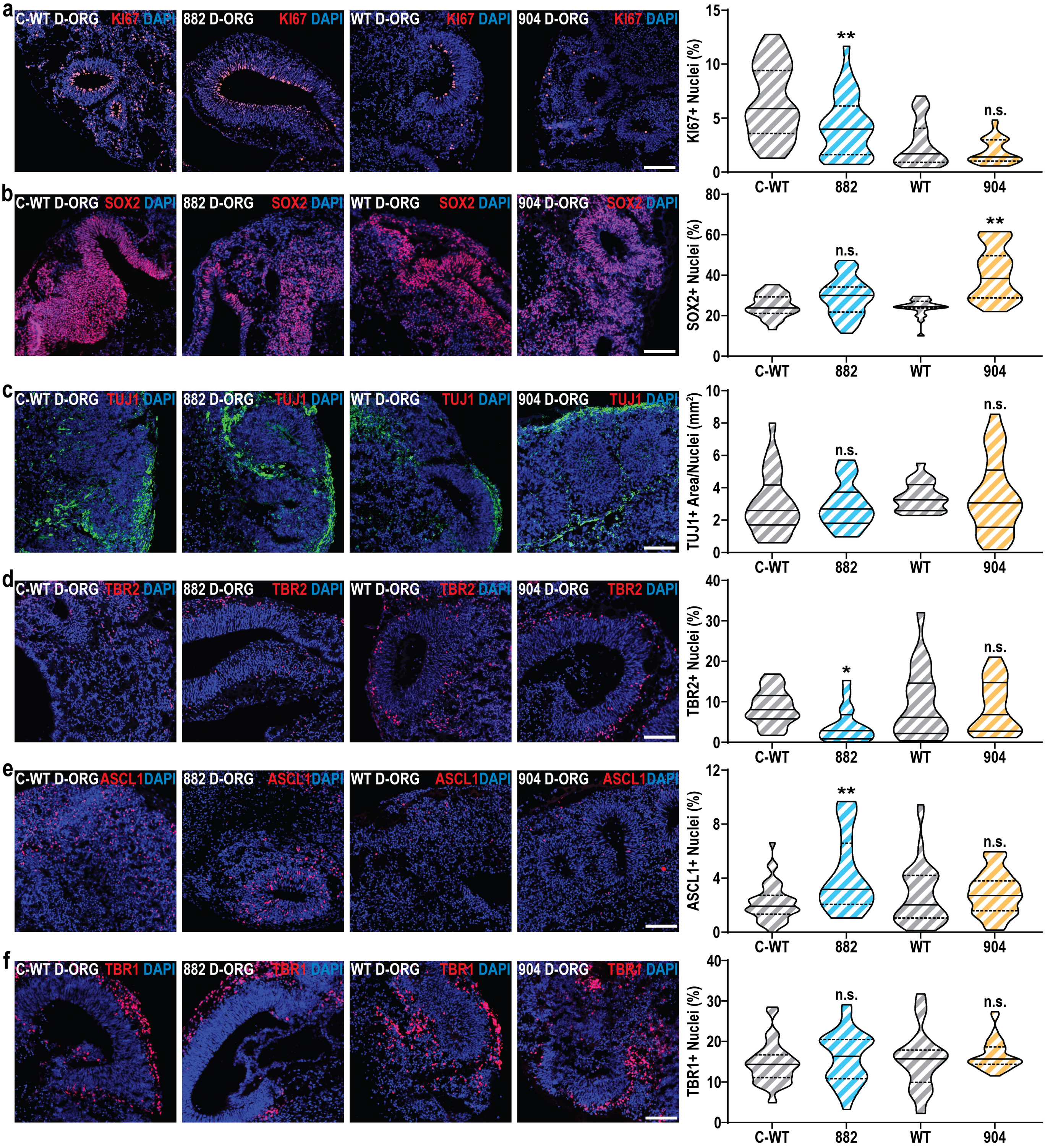


**Supplementary Fig. S7: Changes in proliferation and neurogenesis in TBRS D-ORGs**

**a-b,** Example images and quantification of the proportion of (**a**) KI67+ or (**b**) SOX2+ nuclei in control (C-WT and WT) or TBRS (882 and 904) D-ORGs. **c**, Example images and quantification of the area occupied by TUJ1 positivity per organoid, normalized to the total number of cells within the same organoid, across control and TBRS D-ORGs. **d-f,** Example images and quantification of the proportion of (**d**) TBR2+, (**e**) ASCL1+ or (**f**) TBR1+nuclei in control or TBRS D-ORGs. All data is represented as distributions with median (bold line) and upper and lower quartiles (dotted lines) indicated and was analyzed by Mann-Whitney tests comparing each TBRS model to its appropriate isogenic control. A minimum of 3 batches of organoids were prepared for each condition with 3-9 organoids assessed per batch. pValues: *p<0.05; **p<0.01; n.s.-non-significant. Scale bars=100 μm.

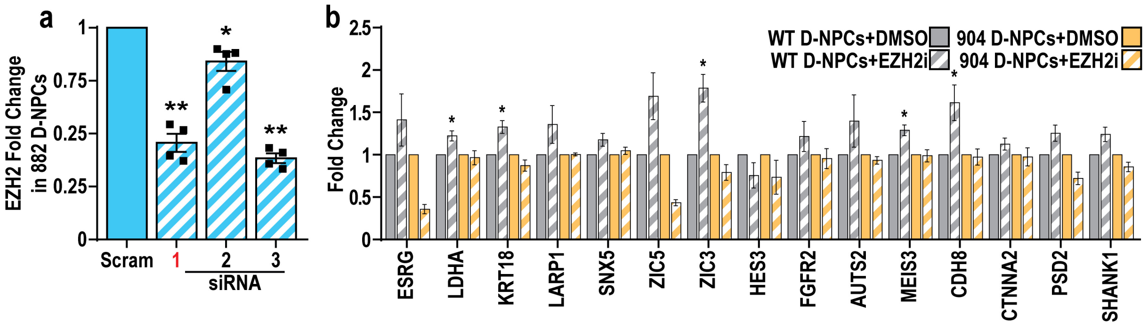


**Supplementary Fig. S8: Consequences of EZH2 modulation in TBRS and control NPCs**

**a,** Quantification of EZH2 mRNA expression in 882 D-NPCs after treatment with a scrambled control siRNA (scram) or siRNAs targeting EZH2 (1-3), highlighting the selection of siRNA 1 (red) for future study. **b,** Quantification of expression changes in 15 genes associated with 882 D-NPC hypo-DMRs and hyper-DKSs in WT and 904 D-NPCs after treatment with an EZH2-specific inhibitor. All data is represented as mean +/- SEM and was analyzed by one-way ANOVA with results shown as changes compared to (**a**) scram or (**b**) DMSO treated conditions. n=4 biological replicate experiments for all conditions, pValues: *p<0.05; **p<0.01 (**a**) or *p<0.05 (**b**).

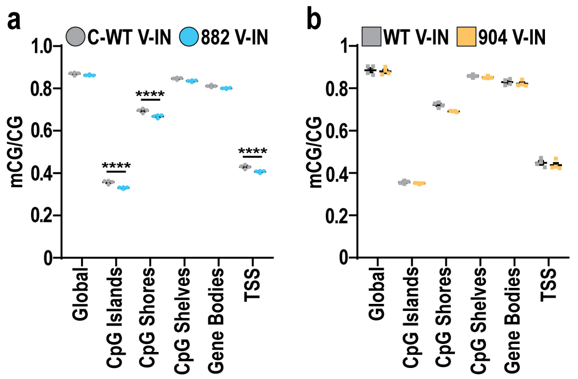


**Supplementary Fig. S9: Changes in DNA methylation in TBRS V-INs**

**a-b,** Changes in mCG/CG levels across (**a**) 882 and (**b**) 904 V-INs compared to matched isogenic control V-INs (C-WT and WT respectively). Data was analyzed by Student's t-test versus isogenic controls (**a-b**). n=4 biological replicate experiments for all conditions, pValues: ****p<0.0001.

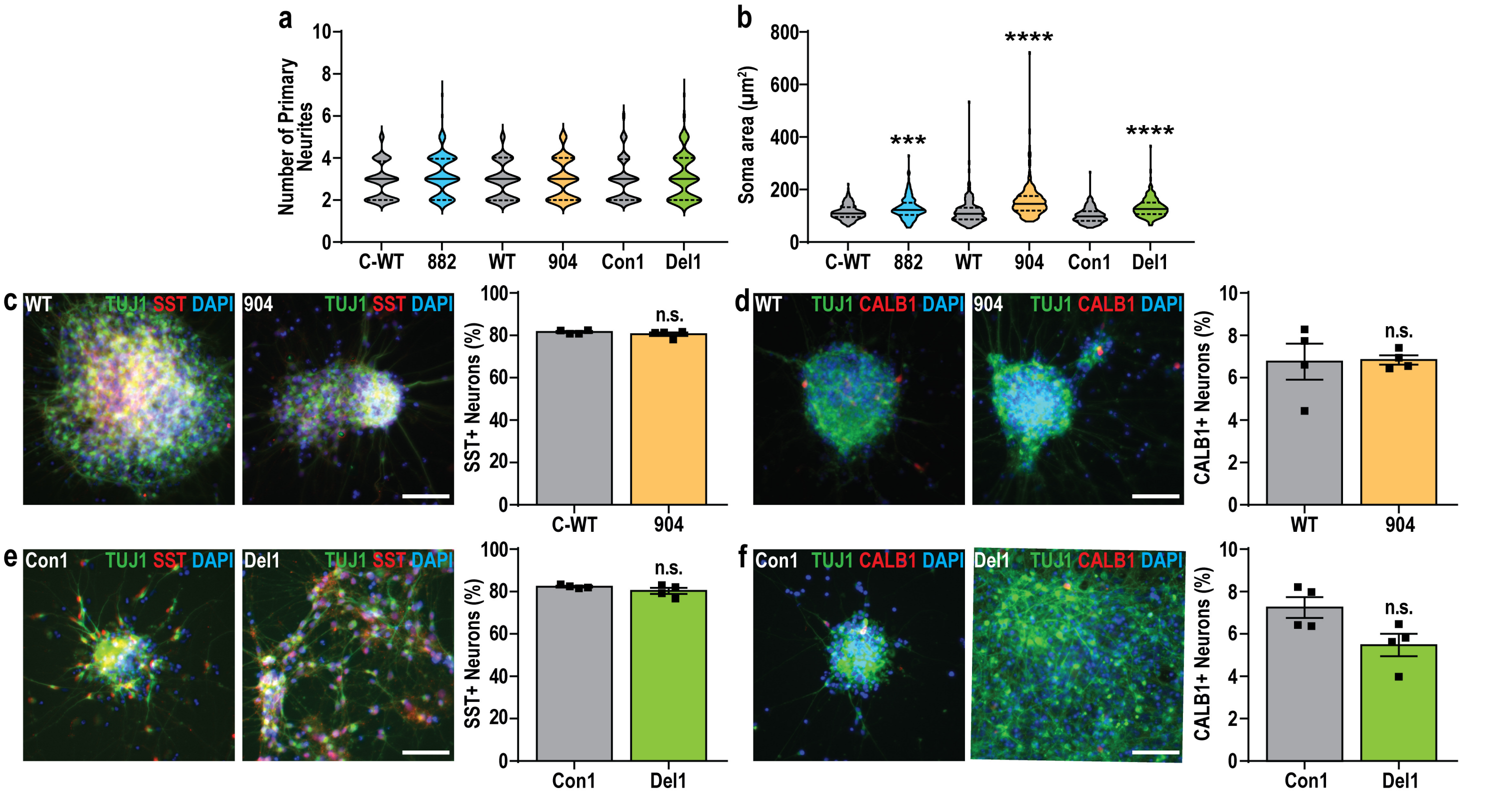


**Supplementary Fig. S10: Assessments of neuronal differentiation in TBRS V-INs**

**a-b,** Quantification of morphological differences between TBRS (882, 904, and Del1) and matched control D30 V-INs, including (**a**) number of primary neurites and (**b**) soma area. **c-f**, Representative images and quantification of the proportion of (c,e) SST+ or (d,f) CALB1+ neurons in (c,d) 904 or (e,f) Del1 D40 V-INs, compared to matched controls (WT and Con1). Data is represented as distributions with median (bold line) and upper and lower quartiles (dotted lines) indicated or as mean +/- SEM. Data was analyzed by Mann-Whitney test (a,b) or Student's t-test (c-f) comparing each TBRS model to its appropriate controls. n=4 biological replicate experiments for all conditions, with a minimum of 20 neurons quantified for morphological measures per biological replicate. pValues: ***p<0.001; ****p<0.0001; n.s.-non-significant. Scale bars=50 µm.

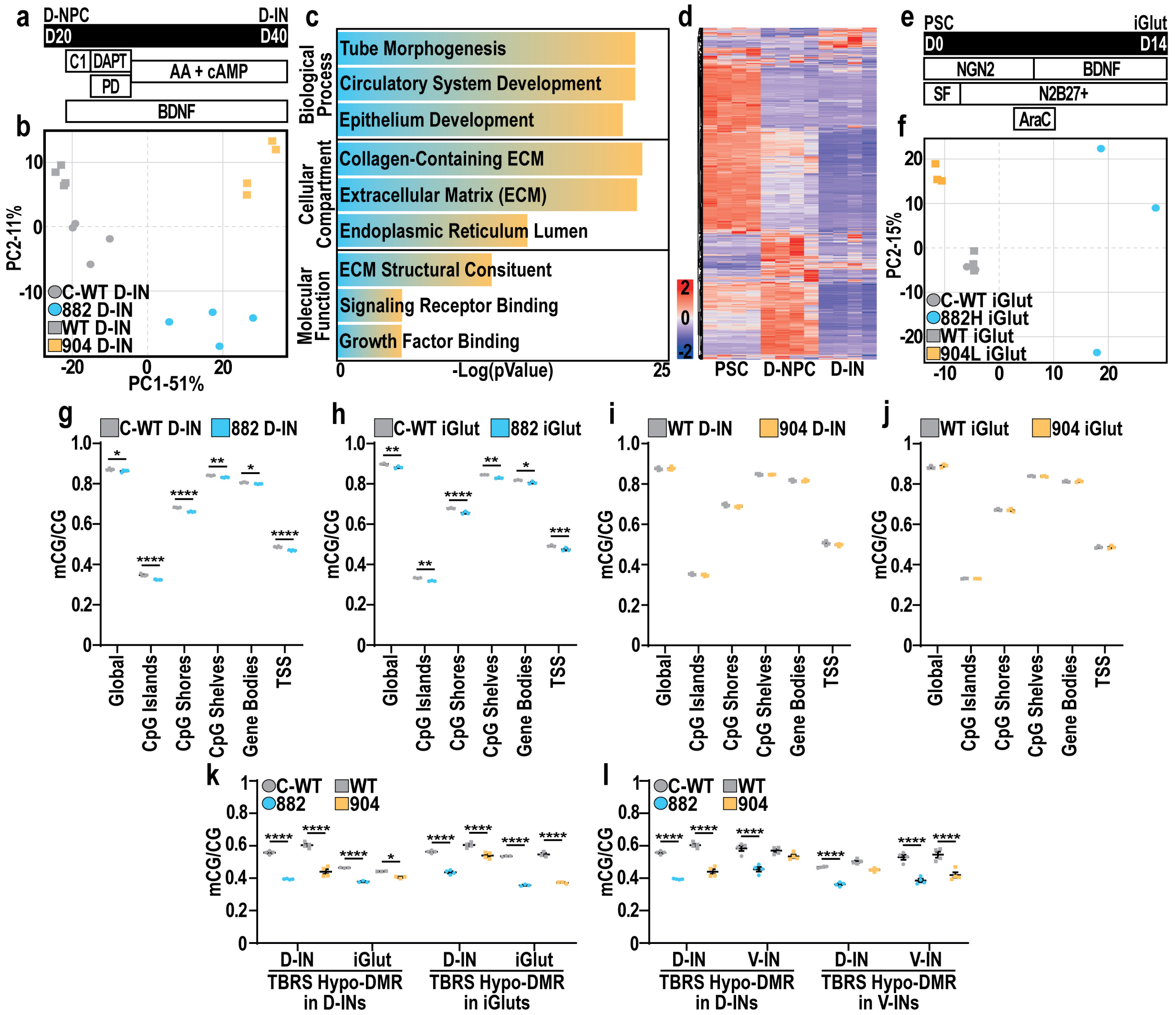


**Supplementary Fig. S11: Characterization of epigenetic and transcriptomic changes in TBRS D-INs**

**a,** Schematic representation of D-IN differentiation from D-NPCs. **b,** Principal component analysis of control (grey) and TBRS D-INs (882-blue and 904-orange). **c,** Summary of GO enrichment analysis for DEGs upregulated in TBRS D-INs. **d**, Heatmap of gene expression from hPSCs to D-INs for DEGs upregulated in TBRS D-INs. **e,** Schematic representation of iGlut differentiation from hPSCs. **f,** Principal component analysis of control (grey) and TBRS iGluts (882-blue and 904-orange). **g-h,** Changes in mCG/CG levels across 882 versus C-WT (**g**) D-INs or iGluts (**h**). **i-j,** Changes in mCG/CG levels across 904 versus WT D-INs (**i**) or iGluts (**j**). **k**, mCG/CG levels at TBRS-shared hypo-DMRs identified in D-INs or iGluts, across both TBRS and control D-INs or iGluts. **l**, mCG/CG levels at TBRS-shared hypo-DMRs identified in D-INs or V-INs, across both TBRS and control D-INs or V-INs. Data was analyzed by Student's t-test versus isogenic controls (**g-l**) . n=4 biological replicate experiments for all conditions, pValues: *p<0.05; **p<0.01; ***p< 0.001; ****p<0.0001.

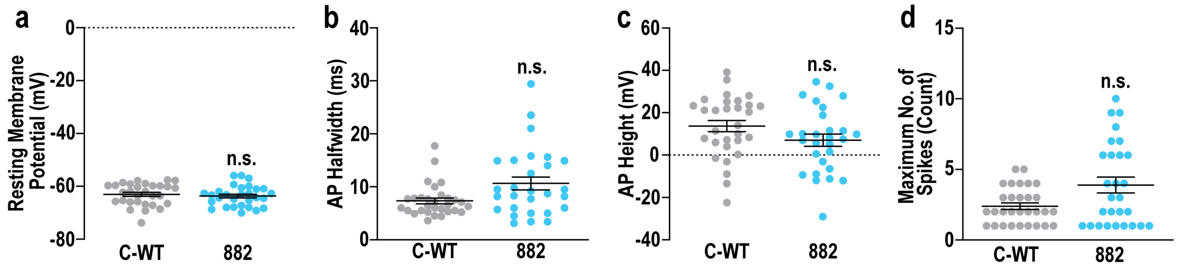


**Supplementary Fig. S12. DNMT3A 882 mutation has no significant effect on glutamatergic neuron function.**
**a-d**, Patch clamp electrophysiology in Day 60 glutamatergic neurons between WT (grey) and 882 (blue) TBRS mutation **a**, Resting membrane potential (mV) demonstrates no significant difference between the electrical potential across the cell membrane under neutral conditions and assessed cells are equally healthy. **b**, Action potential (AP) halfwidth (ms) demonstrating no significant difference between the width of C-WT and 882 glutamatergic action potentials. **c**, Action Potential (AP) height (mV) demonstrating no significant difference between the peak of action potentials of glutamatergic neurons. **d**, Maximum number of spikes (count) during a 1 second depolarizing current pulse, demonstrating no difference in the activity of glutamatergic neurons. Significance was calculated by Rank Sum Test and results are presented as mean +/- SEM, pValues: n.s. p>0.05.

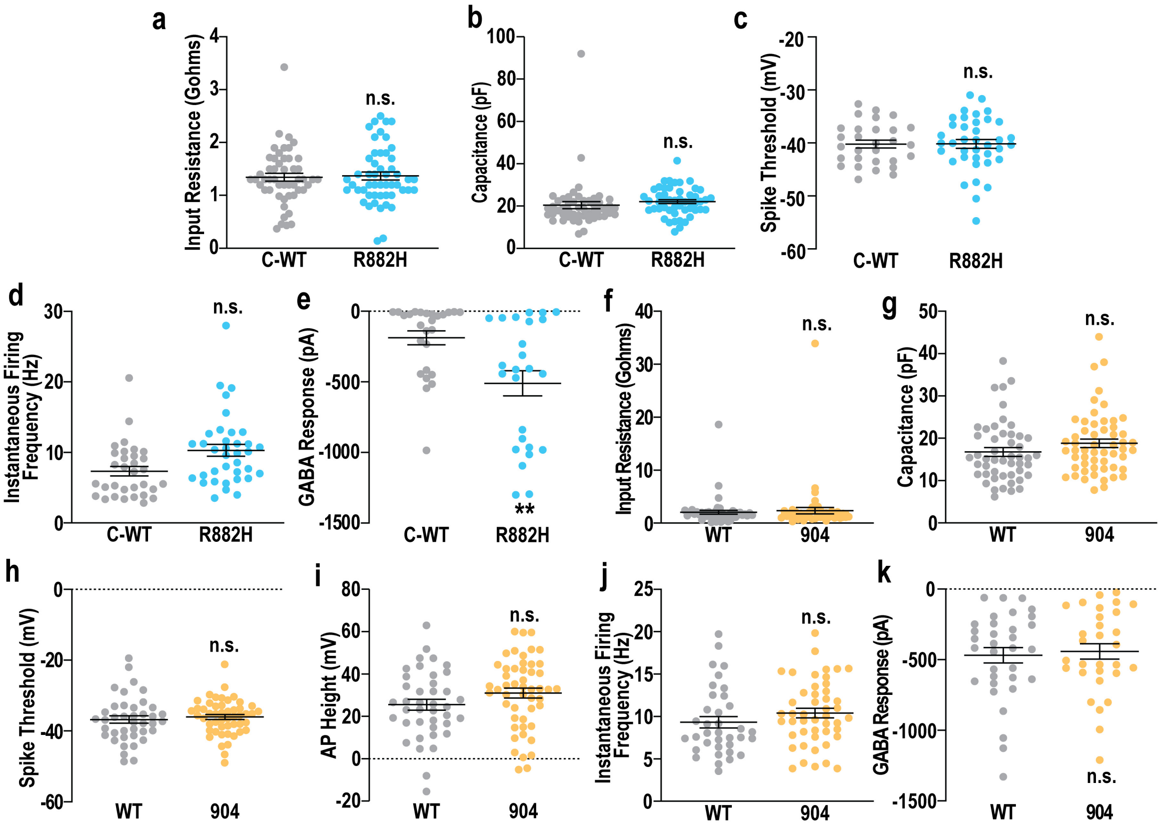


**Supplementary Fig. S13. Unlike the DNMT3A 882 mutation the 904 mutation has no significant affect on GABAergic neuron function.**

**a-e**, Patch clamp electrophysiology in day 60 GABAergic neurons comparing control C-WT (grey) and 882 TBRS (blue) models. **a**, Input resistance (Ω) demonstrating no significant difference between the unstimulated flow of ions and measuring the health of cells. **b**, Capacitance (pF) demonstrating similar surface membrane area. **c**, Spike threshold (mV) demonstrating the cells have similar voltages for action potential initiation. **d**, Instantaneous firing frequency (Hz) demonstrating similar ability for repetitive action potential generation. **e**, GABA Response (pA) demonstrating exposure to 100 µM GABA elicits larger currents in 882 GABAergic neurons compared to C-WT GABAergic neurons. **f-k**, Patch clamp electrophysiology in day 60 GABAergic neurons comparing WT (grey) and 904 (orange) TBRS mutation. **f**, Input resistance (Ω) demonstrating no significant difference between unstimulated flow of ions and measuring the health of cells. **g**, Capacitance (pF) demonstrating similar surface membrane area. **h**, Spike threshold (mV) demonstrating cells have similar voltage for action potential intiation. **i**, Action potential (AP) height (mV) demonstrating no significant difference between the peak of action potentials. **j**, Instantaneous firing frequency (Hz) demonstrating similar ability for repetitive action potential generation. **k**, GABA Response (pA) demonstrating currents elicited by 100 µM GABA are similar in WT and 904 GABAergic neurons. Significance was calculated by Rank Sum Test and results are presented as mean+/-SEM, pValues: n.s. p>0.05,**p<0.01.

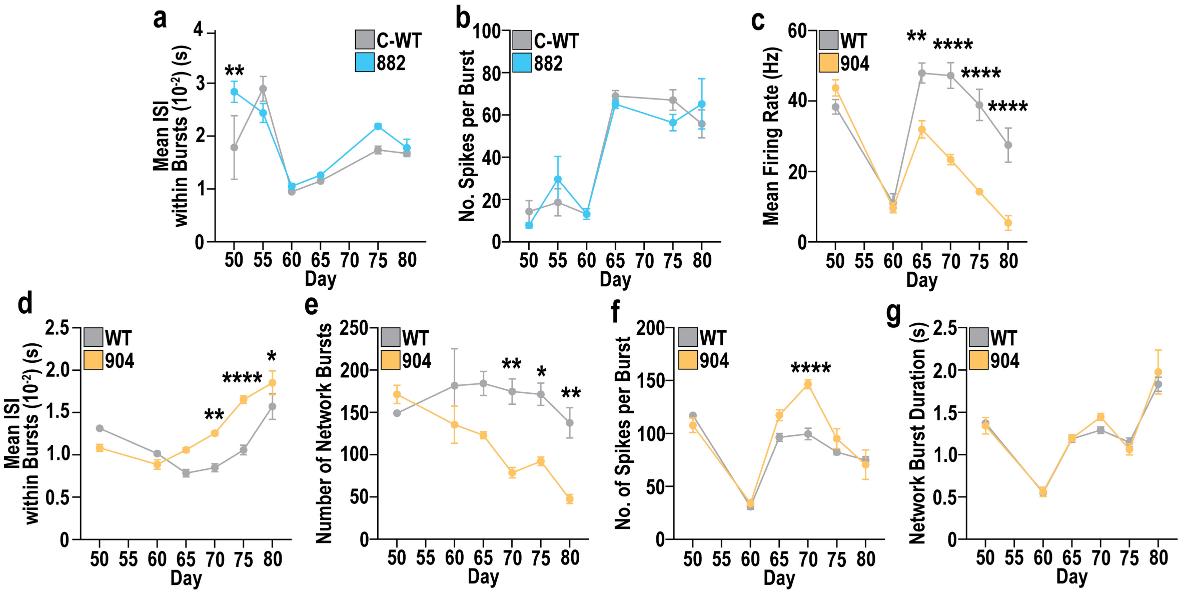


**Supplementary Fig. S14. Low density MEA recordings show decreased activity associated with DNMT3A 904 mutation.**

**a**, Mean inter-spike-interval (ISI) in C-WT (grey) and 882 (blue) glutamatergic neuron and GABAergic neuron mixed cultures recorded from D50-D80 highlights a significant increase in 882 neurons at D50, but no difference across models and differentiation time ($F_{1,6}$=2.55, n=4). **b**, Number of spikes per burst in C-WT and 882 co-cultured neurons demonstrates no significant difference ($F_{1,6}$=0.001, n=4). **c**, Changes in mean firing rate in WT (grey) and 904 (orange) glutamatergic and GABAergic neuron mixed cultures recorded from day (D)50-D80, highlighting a significant decrease in mean firing rate in 904 cultures ($F_{1,6}$=28.2, pValue<0.05, n=4). **d**, Changes in mean inter-spike-interval (ISI) in WT and 904 co-cultures highlighting a significant increase in 904 neurons at days 70-80 and across differentiation time ($F_{1,6}$=13.9, pValue<0.05, n=4). **e**, Changes in the number of network bursts in a 10-minute recording in WT and 904 co-cultured neurons recorded from D50-D80 highlights a significant decrease in the number of network bursts in 904 cultures ($F_{1,6}$=17.4, pValue<0.05, n=4). **f**, Number of spikes per burst in WT and 904 co-cultured neurons, demonstrates a significant increase in 904 neurons at day 70 ($F_{1,6}$=12.23, pValue<0.05, n=4). **g**, Network burst duration in WT and 904 co-cultures recorded from D50-D80 demonstrates no significant difference ($F_{1,6}$=0.48, pValue<0.05, n=4). Significance was calculated by 2-way ANOVA and significant marks represent post-hoc testing with Bonferroni correction for statistical differences between TBRS models and matched isogenic controls at each time point measured, *pValue<0.05; **pValue<0.01 and ****pValue<0.0001.

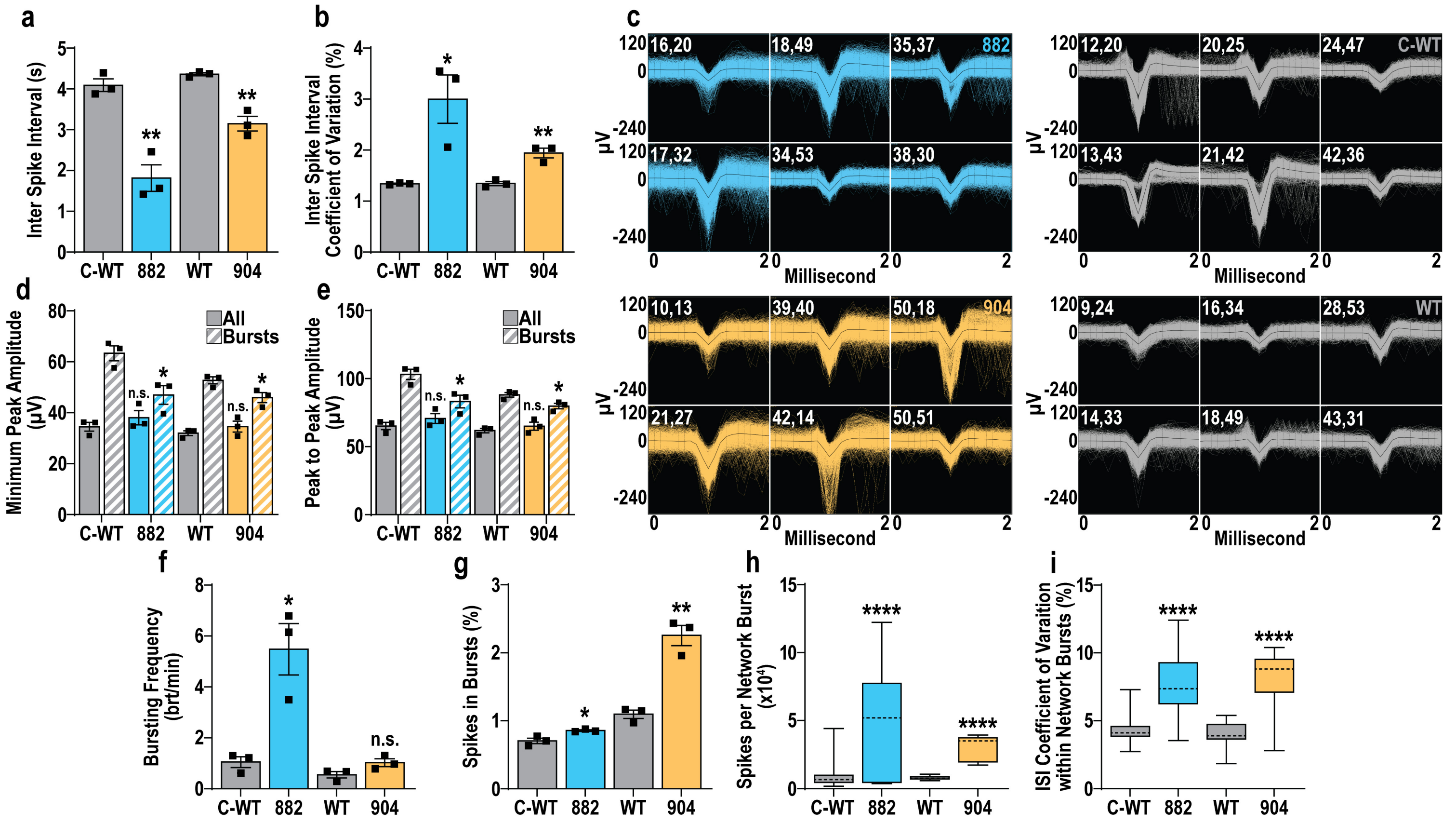


**Supplementary Fig. S15. High density MEA recordings at D82 highlight similar functional and network alterations due to the presence of TBRS GABAergic neurons.**

**a**, Average inter-spike-interval (ISI) of all spikes across cultures of control iGluts and C-WT, WT, 882 and 904 GABAergic neurons at D82 of GABAergic neuron differentiation. **b**, Coefficient of variation (CoV) of ISI of all spikes in cultures with D82 control or TBRS GABAergic neurons. **c**, Example traces of spikes identified in 6 non-contiguous electrodes, showing all spikes identified across a 10-minute recording (blue, grey or orange) and the average spike trace (black) with electrode coordinates marked for each set of traces in cultures with D82 control or TBRS GABAergic neurons. **d**, Minimum peak amplitude for all spikes (solid color) or spikes within electrode bursts (striped color) in cultures with D82 control or TBRS GABAergic neurons. **e**, Peak to peak amplitude for all spikes (solid color) or spikes within electrode bursts (striped color) in cultures with D82 control or TBRS GABAergic neurons. **f**, Electrode bursting frequency in cultures with D82 control or TBRS GABAergic neurons. **g**, Quantifications of the number of spikes in electrode bursts as a proportion of the total number of spikes across 10-minute recordings of cultures with D82 control or TBRS GABAergic neurons. **h**, The average number of spikes per network bursts in cultures with D82 control or TBRS GABAergic neurons. **i**, Quantification of ISI CoV of spikes within network bursts in cultures with D82 control or TBRS GABAergic neurons. Data is represented as mean +/- SEM and was analyzed by Student's t-test, versus matched isogenic controls. n=3 biological replicate experiments for all conditions n.s. pValue>0.05; *pValue<0.05; **pValue<0.01 and ****pValue<0.0001.

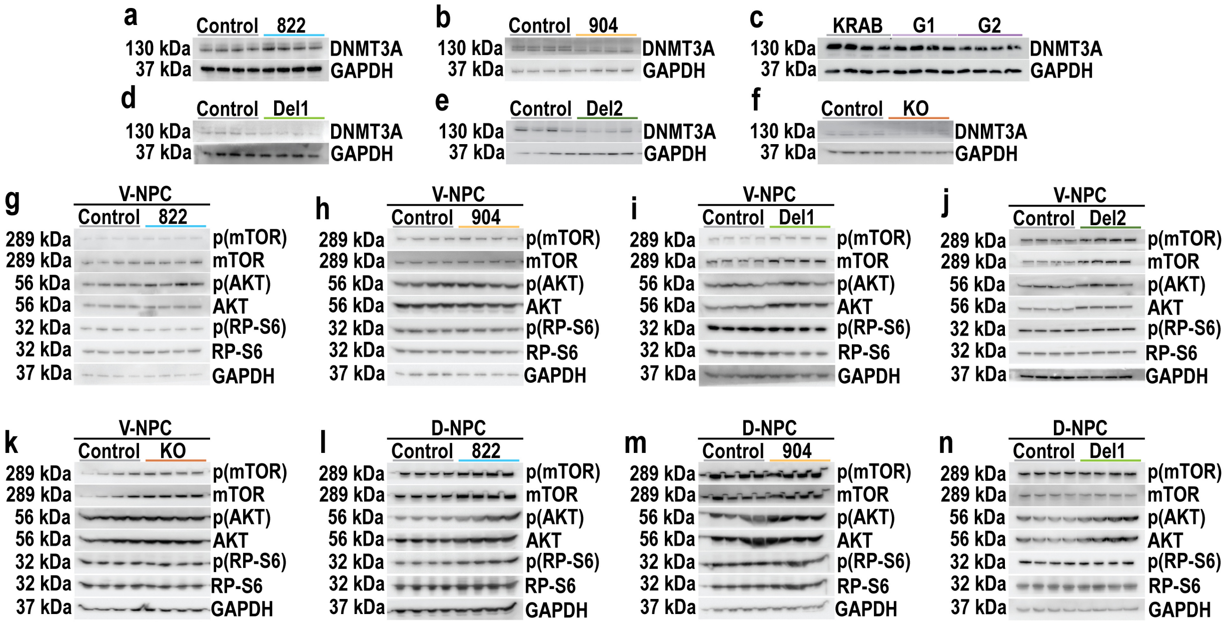


**Supplementary Fig. S16: Complete Western Blots**

**a-f,** Western blots comparing changes in DNMT3A protein levels between matched control and (**a**) 882, (**b**) 904, (**c**) G1/2, (**d**) Del1, (**e**) Del2 and (**f**) KO hPSCs. **g-k**, Western blots of V-NPCs comparing changes in protein levels and phosphorylation of mTOR, AKT and RP-S6 between matched control and (**g**) 882, (**h**) 904, (**i**) Del1, (**j**) Del2 and (**k**) KO. **l-n**, Western blots of D-NPCs comparing changes in protein levels and phosphorylation of mTOR, AKT and RP-S6 between matched control and (**l**) 882, (**m**) 904 and (**n**) Del1.

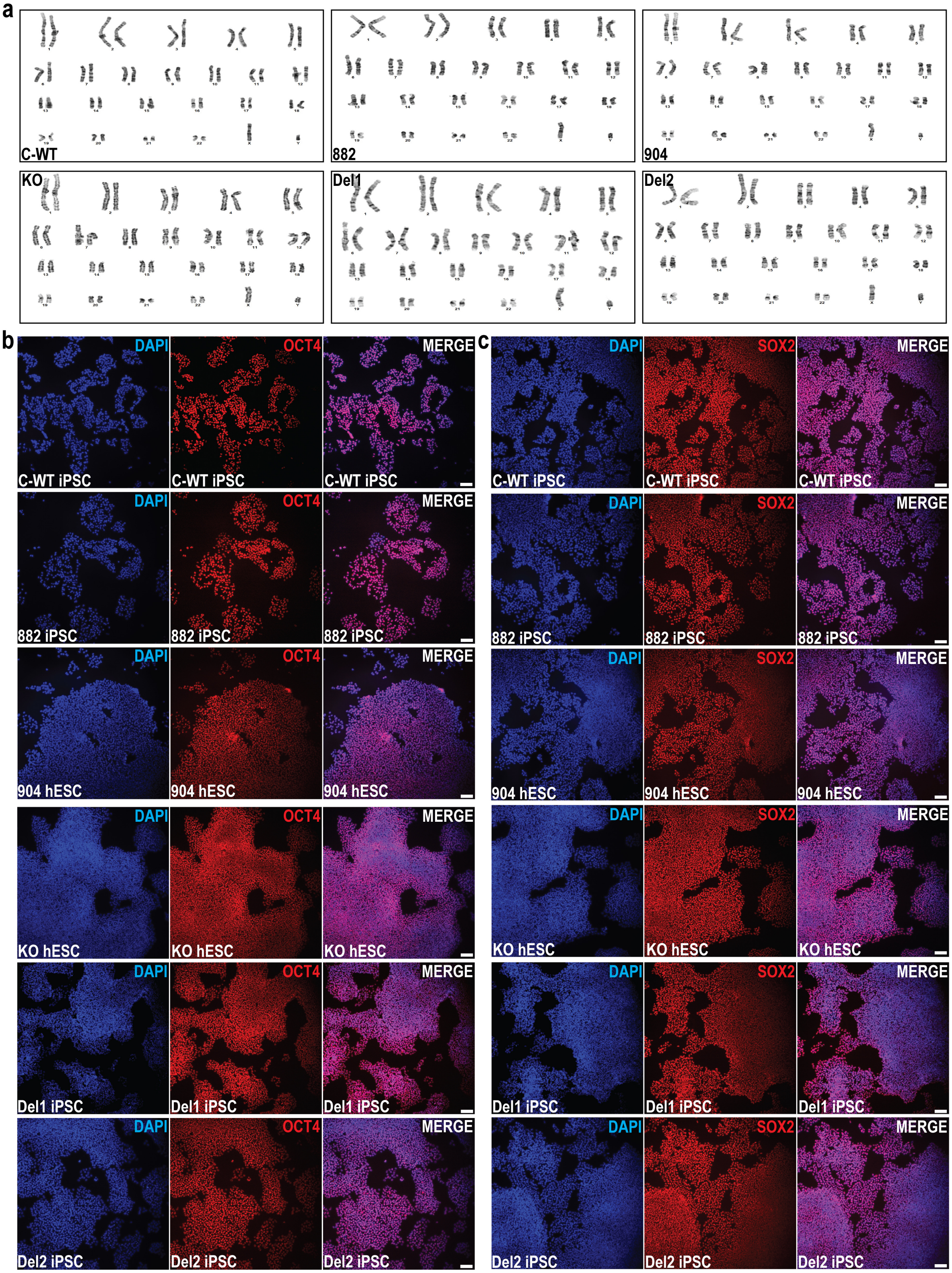


**Supplementary Fig. S17: Characterization and validation of iPSC and hESC lines used in study.**

**(a)** Karyotyping of newly generated iPSC and hESC lines used in study, validating lines are karyotypically normal. **b-c,** Representative images of **(b)** OCT4 and **(c)** SOX2 immunostaining, validating that the hPSC lines used in study express pluripotency markers. Scale bar = 100 µm.

#### References


Chapman G, Determan J, Jetter H, Kaushik K, Prakasam R, Kroll KL. 2024. Defining cis-regulatory elements and transcription factors that control human cortical interneuron development. *iScience* **27**: 109967.

Fantuzzo JA, Mirabella VR, Hamod AH, Hart RP, Zahn JD, Pang ZP. 2017. Intellicount: High-Throughput Quantification of Fluorescent Synaptic Protein Puncta by Machine Learning. *eNeuro* **4**.

Kechin A, Boyarskikh U, Kel A, Filipenko M. 2017. cutPrimers: A New Tool for Accurate Cutting of Primers from Reads of Targeted Next Generation Sequencing. *J Comput Biol* **24**: 1138-1143.

Liu S, Li D, Lyu C, Gontarz PM, Miao B, Madden PAF, Wang T, Zhang B. 2021. AIAP: A Quality Control and Integrative Analysis Package to Improve ATAC-seq Data Analysis. *Genomics Proteomics Bioinformatics* **19**: 641-651.

Love MI, Huber W, Anders S. 2014. Moderated estimation of fold change and dispersion for RNA-seq data with DESeq2. *Genome Biol* **15**: 550.

Meganathan K, Lewis EMA, Gontarz P, Liu S, Stanley EG, Elefanty AG, Huettner JE, Zhang B, Kroll KL. 2017. Regulatory networks specifying cortical interneurons from human embryonic stem cells reveal roles for CHD2 in interneuron development. *Proc Natl Acad Sci U S A* **114**: E11180-E11189.

Patro R, Duggal G, Love MI, Irizarry RA, Kingsford C. 2017. Salmon provides fast and bias-aware quantification of transcript expression. *Nat Methods* **14**: 417-419.

Ramirez F, Ryan DP, Gruning B, Bhardwaj V, Kilpert F, Richter AS, Heyne S, Dundar F, Manke T. 2016. deepTools2: a next generation web server for deep-sequencing data analysis. *Nucleic Acids Res* **44**: W160-165.

Ross-Innes CS, Stark R, Teschendorff AE, Holmes KA, Ali HR, Dunning MJ, Brown GD, Gojis O, Ellis IO, Green AR et al. 2012. Differential oestrogen receptor binding is associated with clinical outcome in breast cancer. *Nature* **481**: 389-393.

Schafer ST, Paquola ACM, Stern S, Gosselin D, Ku M, Pena M, Kuret TJM, Liyanage M, Mansour AA, Jaeger BN et al. 2019. Pathological priming causes developmental gene network heterochronicity in autistic subject-derived neurons. *Nat Neurosci* **22**: 243-255.

Soneson C, Love MI, Robinson MD. 2015. Differential analyses for RNA-seq: transcript-level estimates improve gene-level inferences. *F1000Res* **4**: 1521.

Stirling DR, Swain-Bowden MJ, Lucas AM, Carpenter AE, Cimini BA, Goodman A. 2021. CellProfiler 4: improvements in speed, utility and usability. *BMC Bioinformatics* **22**: 433.
